# Investigating the Impacts of Sphingomyelinases on Extracellular Vesicle Cargo Sorting

**DOI:** 10.1101/2024.12.16.628770

**Authors:** Juan-Carlos A. Padilla, Seda Barutcu, Jonathan Boulais, Yiran Chen, Easin Syed, Eunjeong Kwon, Eric Lécuyer

## Abstract

Extracellular vesicles (EVs) form through regulated biogenesis processes involving sphingomyelinases (SMases), enzymes that metabolize sphingomyelin to produce ceramide—a lipid influencing membrane rigidity and essential for EV generation. This study explores the impact on EV protein and RNA cargoes resulting from inhibiting neutral SMase (NSM) and acid SMase (ASM) in human MCF7 cells. Our results revealed that NSM inhibition reduces EV nanoparticles and diminishes RNA and protein cargoes, including endosomal, spliceosomal, and translation-related proteins. Conversely, ASM inhibition increased RNA-binding proteins within and enhanced the expression of ribonucleoprotein complex-associated RNA in released EVs, including several snRNAs and 7SL RNA. Intriguingly, ASM-inhibited EVs enhanced the migration and translational activity of recipient MCF10A cells. These findings suggest an important role for SMase-dependent vesiculation in governing RNA and protein trafficking to the extracellular space, unveiling potential implications for cellular communication and function.

## INTRODUCTION

Most cells release extracellular vesicles (EVs), heterogeneous membranous nanoparticles, which act as messengers in intercellular communication ^1^. EVs serve as protective shuttles, ferrying diverse bioactive cargoes, including RNAs and RNA-binding proteins (RBPs) ^2–5^. This facilitates their efficient transfer to near and distant microenvironments, ultimately resulting in fusion with recipient cells and influencing their behavior ^6–13^. While the RNA and protein repertoires of EVs are indicative of their parental cells, they are not identical, with EV subpopulations originating from the same cells displaying their own molecular signatures ^14–16^. Unraveling the mechanisms governing this selective packaging is important for understanding the dual roles of EVs as biomarkers and effectors, and further characterization remains imperative.

EV biogenesis is spatially regulated, taking place either locally within the endosomal compartment or at the cell membrane. These processes result in the formation of distinct classes, such as exosomes and microvesicles, respectively ^17,18^. Traversing both EV classes is a metabolic pathway involving sphingomyelinases (SMase), a family of enzymes capable of converting sphingomyelin, a membrane sphingolipid, into ceramide and phosphorylcholine ^19,20^. Neutral sphingomyelinase (NSM) generates exosomes by metabolizing sphingomyelins in the endosomal limiting membrane, forming microdomains that induce negative membrane curvature, ultimately leading to the generation of intraluminal vesicles ^21–24^. Conversely, microvesicles are formed at the plasma membrane by acid sphingomyelinase (ASM), which, upon activation of the ATP receptor P2X_7_, translocates to the outer leaflet of the plasma membrane, where it can influence membrane curvature in a similar fashion to NSM ^25–27^.

Considering the hypothesized distinct contributions of NSM and ASM in EV biogenesis within discrete subcellular compartments, i.e. endosomal networks and the plasma membrane, our study presents a comprehensive comparison of the consequences resulting from inhibiting these enzymatic pathways. Utilizing specific pharmacological inhibitors for NSM (GW4869) ^21,28–30^ and ASM (FTY720/Fingolimod) ^26,31–33^ in human MCF7 adenocarcinoma cells, we systematically evaluated the impact on EV biogenesis, cargo selection, and functional activity. More specifically, by profiling the protein and RNA content of inhibitor-treated MCF7 cells and their secreted EVs, we show that NSM or ASM inhibition leads to distinctive impacts on EV content, particularly concerning ribonucleoprotein machineries, with the former inhibiting and the latter enhancing their secretion within EVs. These findings not only illuminate the divergent roles played by NSM and ASM in EV biogenesis, but also provide a foundation for understanding their contributions to the regulatory landscape of these EVs.

## METHODS

### Cell Culture

Human MCF7 epithelial adenocarcinoma cells were cultured in Minimum Essential Medium MEM 1x (Corning, MT10010CV) supplemented with 10% foetal bovine serum (FBS), 0.1 mM non-essential amino acids (Thermo Fisher, MT25025CI), 1 mM sodium pyruvate (Wisent, 600-110-EL), 100 U mL^-1^ penicillin, and 100 µg mL^-1^ streptomycin (Wisent, 450-202-EL). The cells were incubated at 37°C in a humidified atmosphere with 5% CO_2_. For EV isolations, the same conditions were used, but the media contained 10% EV-depleted foetal bovine serum (dFBS) (EV-isolation media). dFBS was prepared through tangential flow filtration (TFF) using the Labscale TFF System (Millipore-Sigma) equipped with a Pellicon XL50 ultrafiltration module, which had a 300 kDa Ultracell membrane (Millipore-Sigma, PXC300C50).

Human MCF10A non-malignant breast epithelial cells were cultured in a 1:1 mixture of Dulbecco’s Modified Eagle Medium (DMEM) (Wisent, 319-016-CL) and Ham’s F12 Medium (Wisent, 318-010-CL). The media mixture was supplemented with 5% horse serum (HS), 20 ng mL^-1^ human epidermal growth factor (hEGF) (Wisent, 511-110-UM), 500 ng mL^-1^ hydrocortisone (Sigma-Aldrich, H0888-1G), 10 µg mL^-1^ insulin (Gibco, 12585014), 100 ng mL^-1^ cholera toxin from *Vibrio cholerae* (Sigma-Aldrich, C8052-.5MG), 100 U mL^-1^ penicillin and 100 µg mL^-1^ streptomycin (Wisent, 450-202-EL). The cells were incubated at 37°C in a humidified atmosphere with 5% CO_2_.

### Sphingomyelinase Activity Assays

MCF7 cells were plated overnight in complete media at 37°C in a humidified atmosphere with 5% CO_2_. The following day, the media was removed, and the cells were rinsed twice with sterile 1x PBS (Wisent, 311-011-CL). Next, the complete media was supplemented with either SMase inhibitor drugs or vehicle: 0.2% DMSO (Sigma-Aldrich, D8418-100ML), 10 µM GW4869 (Sigma-Aldrich, D1692-5MG), or 5 µM FTY720 (Sigma-Aldrich, SML0700-5MG). The appropriate media was then added to the cells, and they were incubated for 1 h at 37°C in a humidified atmosphere with 5% CO_2_. Following this first incubation, the media was removed, and the cells were rinsed once more with sterile 1x PBS. The treated media was then re-added to the cells, and they were incubated for 36 h at 37°C in a humidified atmosphere with 5% CO_2_. After 36 h, the cells were lysed using Mammalian Cell Lysis Buffer (Abcam, ab179835) according to the manufacturer’s instructions. The protein concentration was determined using the Qubit Protein Assay Kit (Thermo Fisher, Q33211) as recommended. For the quantification of ASM or NSM activity, 200 μg of protein from each condition was used. ASM activity was measured using the Acidic Sphingomyelinase Assay Kit (Fluorimetric) (Abcam, ab190554), while NSM activity was determined using the Sphingomyelin Assay Kit (Fluorometric) (Abcam, ab138877) following the manufacturer’s protocol.

### Treatments with Sphingomyelinase Inhibitors for EV Isolation

MCF7 cells were plated in T-175 flasks in complete media overnight at 37°C in a humidified atmosphere with 5% CO_2_. The following day, the media was removed, and the cells were rinsed twice with sterile 1x PBS (Wisent, 311-011-CL). Next, EV-isolation media was supplemented with either sphingomyelinase inhibitor drugs or vehicle: 0.2% DMSO (Sigma-Aldrich, D8418-100ML), 10 µM GW4869 (Sigma-Aldrich, D1692-5MG), or 5 µM FTY720 (Sigma-Aldrich, SML0700-5MG). The appropriate media was then added to each flask series, and the cells were incubated for 1 h at 37°C in a humidified atmosphere with 5% CO_2_. Following this first incubation, the media was removed, and the cells were rinsed once more with sterile 1x PBS. The treated media was then re-added to each flask series, and the cells were incubated for 36 h at 37°C in a humidified atmosphere with 5% CO_2_. After 36 h, the cell-conditioned media was collected, along with a sample of cells. The cells were assessed for ≥95% cell viability by Trypan blue exclusion assay (Gibco, 15250061). The cell-conditioned media was then used for the purification of EVs.

### EV Isolation by Iodixanol Gradient Fractionation

EVs were purified from the EV-isolation cell-conditioned media based on established methodologies ^34^. In brief, the collected cell-conditioned media underwent centrifugation at 500 × *g* for 10 min and then at 2000 × *g* for 20 min to remove cells and large debris, respectively. The resulting supernatant was collected and concentrated by centrifugation at 4000 × *g* using 100,000 NMWL molecular cutoff Amicon Ultra-15 Centrifugal Filter Units (Millipore-Sigma, UFC910024) to achieve a volume of 1.8 mL. Subsequently, a non-continuous density gradient was generated using OptiPrep (iodixanol) Density Gradient Medium (Sigma-Aldrich, D1556-250ML) with iodixanol concentrations of 5%, 20%, and 30% (sample). The gradient was then subjected to ultracentrifugation for 2.5 h at 200,000 × *g* to separate EVs from other extracellular materials. EV fractions were then collected and diluted in 1x PBS, after which the EVs were pelleted by ultracentrifugation at 150,000 × *g* for 1.5 h. The EV pellets were resuspended in 1x PBS for downstream applications. All steps were conducted at 4 °C.

### EV Isolation by Size Exclusion Chromatography

EVs were purified from the EV-isolation cell-conditioned media by size exclusion chromatography (SEC) with qEVsingle columns (Izon Science, SP2-USD), following the manufacturer’s recommendations. Briefly, the cell-conditioned media was recovered and cleared of cells and large debris by centrifugation at 500 × *g* for 10 min, followed by 2000 × *g* for 20 min. The supernatant was collected and then concentrated to 100 µL by centrifugation at 4000 × *g* using 100,000 NMWL molecular cutoff Amicon Ultra-15 (Millipore-Sigma, UFC910024) and Amicon Ultra-0.5 (Millipore-Sigma, UFC510096) Centrifugal Filter Units. This 100 µL sample was then loaded onto the column and allowed to elute. Next, 900 µL of 0.1 µm-filtered 1x PBS was loaded onto the column and allowed to elute, resulting in a combined 1 mL void volume. Finally, 600 µL of 0.1 µm-filtered 1x PBS was loaded onto the column and recovered as the nanoparticle-enriched fraction.

### Western Blotting

Following the incubation with either vehicle or SMase inhibitor treatments, samples from both cells and EVs were collected. The EVs, which were purified using gradient fractionation, were resuspended in 1x PBS and subsequently lysed using 4x Laemmli sample buffer. In parallel, cellular protein lysates were prepared by sonication. Subsequently, Sodium Dodecyl Sulfate Polyacrylamide Gel Electrophoresis (SDS-PAGE) was carried out using a 10% polyacrylamide gel to separate the proteins based on their molecular weight. For Western blotting, the following primary antibodies were utilized: anti-calnexin (Abcam, ab10286), anti-Tsg101 (GeneTex, GTX118736), and anti-CD81 (GeneTex, GTX101766). The secondary antibody employed was goat anti-rabbit IgG-HRP (Thermo Fisher, 31460).

### Nanoparticle Tracking Analysis

After the application of either vehicle or SMase inhibitor treatments, the EV-isolation cell-conditioned media were recovered and cleared of cells and large debris by centrifugation at 500 × *g* for 10 min, and 2000 × *g* for 20 min respectively. The supernatant was collected and used for Nanoparticle Tracking Analysis (NTA) via the NanoSight NS500 (Malvern Panalytical) system. NTA was performed with the 532 nm laser through three 30-second recordings at 37°C. Data processing and analysis were done using the NanoSight NTA software v3.0 (Malvern Panalytical).

### Transmission Electron Microscopy

EVs, purified through an iodixanol gradient, were visualized using Transmission Electron Microscopy (TEM). This process employed a negative stain approach using uranyl acetate, based on methods previously described ^35^. Briefly, the EV samples were resuspended in 1x PBS, and 5 µL of each sample was loaded onto previously discharged Formvar/carbon-coated copper grids. The samples were allowed to adhere for 3 min. Subsequently, the grids containing the samples were washed three times in droplets of water. This was followed by placement in a 10 µL droplet of 2% uranyl acetate for 1 min. The grids were then blotted to remove excess uranyl acetate and air-dried for 1 h. Finally, the samples were imaged using an FEI Tecnai T12 120kV transmission electron microscope (Field Electron and Ion Company).

### Intra-EV RNA Labeling and Nanoflow Cytometry

After the application of either vehicle or SMase inhibitors, the cell-conditioned media were recovered and cleared of cells and large debris by centrifugation at 500 × *g* for 10 min, followed by 2000 × *g* for 20 min. The supernatant was collected and then concentrated to 50 µL by centrifugation at 4000 × *g* using 100,000 NMWL molecular cutoff Amicon Ultra-15 (Millipore-Sigma, UFC910024) and Amicon Ultra-0.5 (Millipore-Sigma, UFC510096) Centrifugal Filter Units. The 50 µL concentrated media samples were then incubated with 50 µL of 0.1 µm-filtered 1x PBS containing 100 µM Syto RNASelect (Thermo Fisher, S32703) (final concentration 50 µM) at room temperature for 2 h, protected from light. Next, the samples were loaded onto qEVsingle columns (Izon Science, SP2-USD) for the purification of EVs and the removal of unbound dye. The stained EVs were then used for nanoflow cytometry experiments.

Nanoflow cytometry of EVs was carried out using a CytoFLEX flow cytometer (Beckman-Coulter) equipped with violet (405 nm), blue (488 nm) and red (640 nm) wavelength lasers. Acquisition and analysis of data was carried out with the CytExpert 2.4 software (Beckman-Coulter) using the parameters detailed on **Table 1**. The violet side scatter (V-SSC) was performed using the 405 nm laser along with a gain of 100 on V-SSC and a threshold of 950 on the violet channel. The samples were loaded to a volume of 10 µL, with a slow sample flow rate of 10 µL min^-1^. Background signal was calculated by acquiring 0.1 µm-filtered 1x PBS (EV suspension buffer) at the same rate.

**Table 1:** Nanoflow Cytometry Parameters Using CytoFLEX.

|  |  |  |
| --- | --- | --- |
| Volume to Record | 10 $\mu\text{L}$ | |
| Sample Flow Rate | 10 $\mu\text{L min}^{-1}$ | |
| Acquisition Setting | Gain |  |
|  | FSC | 22 |
|  | SSC | 254 |
|  | Violet SSC | 100 |
|  | FITC | 3000 |
|  | APC | 3000 |
|  | Threshold |  |
|  | Channel | Violet |
|  | Manual, 950>0 (Height) |  |

### Protein Preparation for Mass Spectrometry

After incubation with either vehicle or SMase inhibitor treatments, samples from both cells and EVs were collected. The EVs were purified using gradient fractionation, and both the EVs and their cells of origin were resuspended in TRIzol reagent (Thermo Fisher, 15596018). This allowed for simultaneous isolation of proteins and RNA from the same samples, following the manufacturer’s recommendations. The cellular and EV samples were homogenized in 500 µL of TRIzol reagent. Then, 100 µL of chloroform (VWR, BDH1109-4LG) was added and the samples were centrifuged at 12,000 × *g* for 15 min. After centrifugation, the aqueous phase was saved for RNA isolation. The interphase and organic phase were processed for protein precipitation as described by the manufacturer. For the EV samples, protein extracts were re-solubilized in 20 µL of 8 M urea. This was achieved by vortexing on a MixMate (Eppendorf) at 1200 RPM for 10 minutes and sonication in a water bath for 10 minutes. For the whole cell protein extracts (WC), 100 µL of 8 M urea, 50 µL of 0.1% SDS and 50 µL of acetonitrile were added to each sample. The same procedure as the EV samples was used to re-solubilize WC samples. Proteins were reduced by adding 5.0 µL (EV) or 50 µL (WC) of the reduction buffer (45 mM DTT, 100 mM ammonium bicarbonate) for 30 min at 37°C. They were then alkylated by adding 5.0 µL (EV) or 50 µL (WC) of the alkylation buffer (100 mM iodoacetamide, 100 mM ammonium bicarbonate) for 20 min at 24°C in the dark. Before trypsin digestion, 75 µL of 50 mM ammonium bicarbonate was added to EV samples to reduce the urea concentration to under 2 M for the digestion step. For the WC samples, 175 µL of 50 mM ammonium bicarbonate was added to reduce the urea concentration to under 2 M and also to reduce the concentration of SDS (0.01%) and acetonitrile (10%). 20 µL and 25 µL of the trypsin solution (100 ng µL^-1^ of trypsin sequencing grade from Promega) were added to EV and WC samples, respectively. Protein digestion was performed at 37°C for 18 h. Protein digests were dried down in a vacuum centrifuge and stored at -20°C.

### Liquid Chromatography Tandem Mass Spectrometry

Before liquid chromatography tandem mass spectrometry (LC-MS/MS), two different sample cleaning methods were used. EV protein digests were re-solubilized under agitation for 15 minutes in 10 µL of 0.2% formic acid. Desalting/cleanup of the digests was performed using C18 ZipTip pipette tips (Millipore). Eluates were dried down in a vacuum centrifuge and stored at -20 °C until LC-MS/MS analysis. WC protein digests were acidified with trifluoroacetic acid for desalting and removal of residual detergents by MCX (Waters Oasis MCX 96-well Elution Plate) following the manufacturer’s instructions. After elution in 10% ammonium hydroxide /90% methanol (v/v), samples were dried down in a vacuum centrifuge and stored at -20 °C until LC-MS/MS analysis. EV and WC samples were reconstituted under agitation for 15 minutes in 15 µL and 120 µL of 2% ANC-1% FA, respectively, and between 2 and 6 µL were loaded into a 75 μm i.d. × 150 mm Self-Pack C18 column installed in the Easy-nLC II system (Proxeon Biosystems). The buffers used for chromatography were 0.2% formic acid (buffer A) and 100% acetonitrile/0.2% formic acid (buffer B). Peptides were eluted with a two-slope gradient at a flow rate of 250 nL min^-1^. Solvent B first increased from 2 to 35% in 124 minutes and then from 35 to 85% B in 12 minutes. The HPLC system was coupled to an Orbitrap Fusion mass spectrometer (Thermo Scientific) through a Nanospray Flex Ion Source. Nanospray and S-lens voltages were set to 1.3-1.8 kV and 60 V, respectively. The capillary temperature was set to 250 °C. Full scan MS survey spectra (m/z 360-1560) in profile mode were acquired in the Orbitrap with a resolution of 120,000 with a target value at 3e5. The 25 most intense peptide ions were fragmented in the HCD collision cell and analyzed in the linear ion trap with a target value at 2e4 and a normalized collision energy at 29. Target ions selected for fragmentation were dynamically excluded for 30 seconds after two MS/MS events.

### Proteomics Bioinformatics Analyses

LC-MS/MS .RAW data were analyzed with MaxQuant software (version 1.6.17.0). MS/MS spectra were searched against the human proteome of the UniProt database (July 10, 2020 release) supplemented with MaxQuant’s contaminants option. ITMS match tolerance was set at 0.5 Da, ITMS de novo tolerance was set at 0.25 Da, while deisotoping tolerance was set at 0.15 Da. A maximum of 2 missed cleavages by trypsin was allowed with a maximum of 5 modifications per peptide. Carbamidomethylation of cysteine residues was set as a fixed modification, whereas methionine oxidation was set as variable modifications. False-discovery rate (FDR) for peptides and proteins was set at 1%, with a minimum peptide length of 7, and a minimum of 1 unique peptide was required. Match between runs was enabled with a match time window of 2.5 minutes and an alignment time window of 20 minutes. Cellular and EV .RAW data were analyzed separately. MaxQuant results were then analyzed in R (www.r-project.org) with the MSstats package ^36^. Once imported to the MSstats format, the data were processed by MSstats with the “highQuality” featureSubset, min_feature_count set at 2, while remaining parameters were set as default. Statistical contrasts were also performed by MSstats with the groupComparison function. Statistically significant proteins (adjusted p-value ≤ 0.05 and Log_2_ fold change ≤ -1 or ≥ 1) from extracellular vesicles contrasts were selected, and each protein was Z-scored separately among EV and WC samples using mean protein intensities. Unidentified proteins among triplicate conditions were considered as biologically missing and accordingly assigned a mean intensity value of 0. With the ComplexHeatmap R package ^37^, a heatmap was generated by performing a first round of clustering with the “complete” agglomeration method on EV samples with Euclidean distances. After extracting 6 EV clusters, each cluster of proteins was individually re-clustered using the “complete” agglomeration method on EV and WC samples on Euclidean distances. Pearson correlations were calculated with mean protein intensities in R with the stats package and plotted with the corrplot package, whereas MDS (Multidimensional scaling) analyses were performed with the limma package ^38^. Overrepresentation analyses (ORA) of GO-BP (biological processes) were executed on the g:Profiler website (<u>biit.cs.ut.ee/gprofiler/gost</u>) on statistically significant EV_FTY720 or EV_GW4869 proteins (adjusted p-value ≤ 0.05 and Log_2_ fold change ≤ -1 or ≥ 1) while applying the Benjamini-Hochberg FDR correction method. A network-based visualization and simplification of statistically significant GO-BP (adjusted p-value ≤ 0.05) were next performed. GO-BP with term sizes of less than 500 genes were loaded in Cytoscape’s EnrichmentMap application ^39^ where similar GO-BP were organized in clusters. Each Cluster was next simplified by assigning a resuming GO-BP in Adobe Illustrator (Adobe Inc.). RNA-binding proteins (RBPs) were identified based on a list obtained from the Encyclopedia of RNA Elements (ENCORE) consortium (https://www.encodeproject.org/), consisting of 1072 proteins. GraphPad Prism (GraphPad Software Inc.) was used to generate volcano plots of differentially expressed proteins in cells, as well as sub-heatmaps of peptide intensities (relative Z-score).

### RNA Preparation for RNA Sequencing

After incubation with either vehicle or SMase inhibitors, samples from both cells and EVs were collected. The EVs were purified using gradient fractionation, and both the EVs and their cells of origin were resuspended in TRIzol reagent (Thermo Fisher, 15596018). This allowed for simultaneous isolation of proteins and RNA from the same samples, following the manufacturer’s recommendations. The cellular and EV samples were homogenized in 500 µL of TRIzol reagent. Then, 100 µL of chloroform (VWR, BDH1109-4LG) was added, and the samples were centrifuged at 12,000 × *g* for 15 min. After centrifugation, the aqueous phase was processed for RNA isolation, while the interphase and organic phase were saved for protein precipitation. Following the centrifugation step, the aqueous phase was processed for RNA purification using the RNA Clean & Concentrator-5 system (Zymo Research, R1013) and in-column DNaseI treatment. All steps were performed according to the manufacturers’ protocols. The RNA was eluted in nuclease-free water and quantified by Nanodrop and Bioanalyzer. After the extraction of RNAs, the samples were ribo-depleted followed by library preparation with the KAPA RNA HyperPrep Kit with RiboErase (HMR) (Roche, 8098140702). Next, starting with 1 µg of total RNA for cellular samples or 2-6 ng of total RNA recovered from EV samples from drug-induced conditions, the libraries were amplified for 7 and 16 PCR cycles, respectively. The libraries were quantified by qPCR and loaded at equimolar concentrations for sequencing on the Novaseq platform with the kit NovaseqS4 at a coverage of 35M paired-end reads per sample.

### Transcriptomics Bioinformatics Analyses

FASTQ sequencing files were assessed for quality control with FastQC (The Babraham Institute: https://www.bioinformatics.babraham.ac.uk/projects/fastqc/). The generated RNA sequencing libraries were mapped to the human reference genome (GRCh38), with GENCODE v34 gene annotation, using the STAR alignment tool ^40^. The mapped reads per gene were counted using featureCounts ^41^. Genes that have a max read number value less than 10 are filtered out prior to pairwise comparison of the samples. Genes differentially enriched in EV and cell lysate samples were determined by pairwise comparison using DESeq2 ^42,43^ with parameters: Fit Type=parametric, betaPrior=FALSE. We used a threshold of Log_2_ fold change (LFC) ≥ 1 or ≤ -1 and adjusted p-value ≤ 0.05 to determine significant hits. The significant hits were further assessed for gene ontology (GO) enrichment analysis using the FuncAssociate software v3.0 ^44^. Pheatmap and UpSetR R packages were used to plot the heatmaps and upset plots respectively.

### Scratch-Wound Wound Healing Assay

MCF10A cells were plated at 20,000 cells per well of a Incucyte Imagelock 96-well plate (Sartorius, BA-04857) and incubated until confluent at 37°C in a humidified 5% CO_2_ atmosphere. Once confluent, and directly before addition of MCF7 EVs, the MCF10A cells were wounded with the Incucyte 96-well WoundMaker Tool (Essen Bioscience, 4563) according to the manufacturer’s recommended protocol.

EVs were purified from MCF7 cells under either vehicle or SMase inhibitors. Briefly, 1.2×10^7^ MCF7 cells were plated overnight in a T-175 flask for each condition. On the following day, the plating media was removed and replaced with EV-isolation media containing vehicle or EV-inhibitor drugs, and the cells were incubated for 1 h at 37°C in a humidified atmosphere with 5% CO_2_. Following this first incubation, the media was removed, and the cells were rinsed once more with sterile 1x PBS. The treated media was then re-added to each flask series, and the cells were incubated for 36 h at 37°C in a humidified atmosphere with 5% CO_2_. The EV-isolation cell-conditioned media were recovered and processed for the isolation of EVs by SEC. Purified EVs were recovered in equal volumes of 1x PBS, and volumetric equivalents were added to wounded MCF10A cells. An equal volume of 1x PBS was also added as a second control. The cells were then live imaged hourly over 48 h with an Incucyte Microscope (Sartorius) to evaluate the effects of co-incubation with EVs. The Incucyte software Scratch Wound Analysis Software Module (Sartorius) was used to calculate the percentage of Relative Wound Density (%RWD), which measures the spatial cell density in the wound area relative to the spatial cell density outside of the wound area at every time point.

### Cell Proliferation Assay

MCF10A cells were wounded as in the scratch-wound healing assay and then incubated at 37°C in a humidified 5% CO_2_ atmosphere with either MCF7 control or inhibitor EVs. Before incubation, the purified EVs in 1x PBS were mixed with 5-ethynyl-2’-deoxyuridine (EdU), a component of the Click-iT EdU Cell Proliferation Kit for Imaging (Invitrogen, C10337), to a final concentration of 10 µM and administered to MCF10A cells. An equal volume of blank 1x PBS with EdU was also added as a second control. The incubation period proceeded for up to 3, 6, 18, or 24 hours, after which the MCF10A cells were fixed with a solution of freshly prepared 3.7% formaldehyde (Fisher Chemical, Cat. No. F79-1) in 1x PBS and labeled by Click chemistry for imaging as per the manufacturer’s recommended protocol. Image acquisition of fluorescently labeled cells was carried out with the ImageXpress Micro High-Content Screening (HCS) Microscopy system (Molecular Devices, LLC), and image analysis was performed using the MetaXpress 3.1 software (Molecular Devices, LLC). Statistical analysis was conducted using GraphPad Prism (GraphPad Software Inc.).

### Protein Synthesis Assay

MCF10A cells were wounded as in the scratch-wound healing assay and then incubated at 37°C in a humidified 5% CO_2_ atmosphere with either MCF7 control or SMase inhibitor EVs. The incubation period proceeded for up to 3, 6, 18, or 24 h, after which the media was replaced with fresh EV-isolation media containing 20 µM O-propargyl-puromycin (OPP), a component in the Click-iT Plus OPP Protein Synthesis Assay Kit (Molecular Probes, C10456), and further incubated at 37°C in a humidified 5% CO_2_ atmosphere for an additional 30 min. The cells were then fixed with a solution of freshly prepared 3.7% formaldehyde (Fisher Chemical, Cat. No. F79-1) in 1x PBS and labeled by Click chemistry for imaging according to the manufacturer’s recommended protocol. Image acquisition of fluorescently labeled cells was performed with the ImageXpress Micro High-Content Screening (HCS) Microscopy system (Molecular Devices, LLC), and image analysis was carried out using the MetaXpress 3.1 software (Molecular Devices). Statistical analysis was conducted using GraphPad Prism (GraphPad Software Inc.).

## RESULTS

### Sphingolipid Metabolism Influences the Release of EV RNA from Cells

To assess the influence of SMases on EV biogenesis and molecular composition, we aimed to disrupt these pathways using pharmacological approaches in human MCF7 breast adenocarcinoma cells. To achieve this, we employed GW4869, a cell-permeable non-competitive inhibitor of neutral sphingomyelinase (NSM), known to disturb endosomal transport by reducing intraluminal vesicle (ILV) formation ^21,30^. Additionally, we utilized FTY720 (Fingolimod), which has been documented to hinder microvesicle formation at the plasma membrane by making acid sphingomyelinase (ASM) susceptible to protease degradation ^26,31^. Following titration experiments, we identified doses of GW4869 (10 µM) and FTY720 (5 µM) that showed no discernible impact on MCF7 cell viability (**Figure 1A**), or proliferation (**Figure S1A**) compared to control cells after 36 hours—the designated time for collecting EVs from the cell-conditioned media. These precautions were taken to avoid contamination of purified populations with apoptotic bodies or effects on the EV population caused by changes in cell numbers.

**FIGURE 1:**
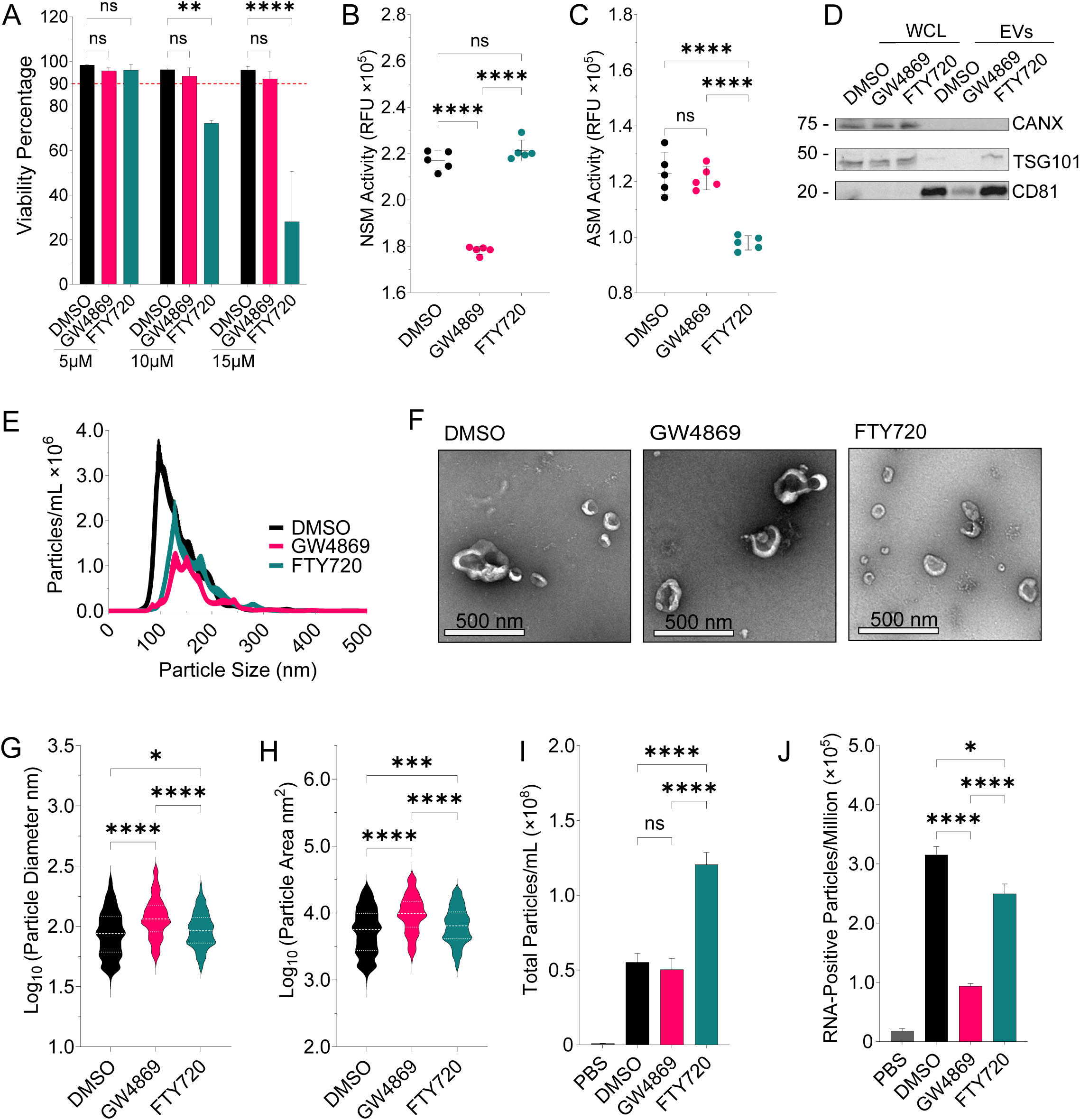
Sphingolipid Metabolism Influences the Release of EV RNA from Cells. **A.** Cellular viability of MCF7 cells was measured after a 36-hour incubation with sphingomyelinase (SMase) inhibitor drugs at different doses. Appropriate dosing concentrations were determined as 5 μM for FTY720 (ASM inhibitor) and 10 μM for GW4869 (NSM inhibitor), with DMSO (vehicle) used as the control condition in a volumetric equivalent volume of 0.2% relative to the total media. All subsequent experiments utilize these doses. **B.** Activity of neutral sphingomyelinase (NSM) as quantified by measuring of relative fluorescence units (RFU) in cellular lysates derived from control or SMase inhibitor-treated cells. **C.** Activity of acid sphingomyelinase (ASM) as quantified by measuring of relative fluorescence units (RFU) in cellular lysates derived from control or SMase inhibitor-treated cells. **D.** Western blot of cellular and EV proteins isolated from control cells or cells treated with SMase inhibitor drugs. The samples were probed for EV markers TSG101 and CD81, as well as the negative control marker calnexin (CANX). **E.** Nanoparticle Tracking Analysis (NTA) performed on 0.45 μm filtered cell-conditioned media collected after incubation of either control cells or cells treated with SMase inhibitor drugs. The standard error is represented by the thickness of the border. **F.** Transmission electron micrographs (TEM) depicting iodixanol gradient-purified EVs derived from either control cells or cells treated with SMase inhibitor drugs. The EVs were negatively stained with 2% uranyl acetate and imaged using a FEI Tecnai T12 120kV transmission electron microscope. **G.** Nanoparticle diameter distributions, measured in nanometers (nm), as quantified using TEMs of EVs with the TEM ExosomeAnalyser tool. **H.** Distribution of total particle area, measured in square nanometers (nm^2^), as quantified using TEMs of EVs with the TEM ExosomeAnalyser tool. **I.** Quantifications of total detected nanoparticles, isolated via size exclusion chromatography (SEC). Measurements were recorded using nanoflow cytometry on the CytoFLEX platform. **J.** Quantifications of RNA-positive nanoparticles, isolated via SEC. Measurements were recorded using nanoflow cytometry on the CytoFLEX platform. RNA labelling was done using SYTO RNASelect, a cell-permeable stain for nucleic acids that is selective for RNA. For all graphs, unless otherwise stated, the error bars represent the standard error of the mean (SEM) and the statistics were calculated by ordinary one-way ANOVA with Tukey post-hoc test (where ns=*P*>0.05, *=*P*≤0.05, **=*P*≤0.01, ***=*P*≤0.001, and ****= *P*≤0.0001) on GraphPad Prism.

Next, we aimed to validate the impact of the pharmacological compounds on the activity of sphingomyelinase enzymes in treated cells. Our results confirmed that the incubation of MCF7 cells with GW4869 or FTY720 at the selected doses led to a specific reduction in NSM and ASM activity in cell lysates (**Figure 1B-C**). To begin characterizing our EV specimens, we performed Western blots of cellular and EV extracts, which revealed an enrichment of well-established EV markers, including TSG101 and CD81, coupled with the depletion of the control marker calnexin (CANX). Interestingly, while EV markers were detected across the control or inhibitor EV specimens, there were noticeable differences in their quantities across equal volumes of purified EVs (**Figure 1D**).

Nanoparticle Tracking Analysis (NTA) of cell-conditioned media unveiled alterations in the size distribution of nanoparticles (**Figure 1E**), indicating an increase in the median size of nanoparticles in GW4869 (162.6 ± 5.0 nm) and FTY720 (161.6 ± 2.6 nm) samples compared to DMSO-treated controls (135 ± 3.3 nm). Examination of purified EV specimens by transmission electron microscopy (TEM) revealed the presence of EVs of diverse sizes under all conditions (**Figure 1F**), with quantifications showing that NSM- and ASM-inhibited samples displayed nanoparticle populations with larger diameters and mean particle areas (**Figure 1G-H**). Interestingly, nanoflow cytometry analysis showed specific effects of the inhibitors on EV populations (**Figure 1I-J**, **S1B**). There was an increase in the total number of particles detected per milliliter in the ASM-inhibited (FTY720) EV population (**Figure 1I**). However, there was an overall reduction in the number of RNA-positive particles (per million events) resulting from both NSM and ASM inhibitor treatments (**Figure 1J**), with GW4869 treatments showing the most pronounced reduction in RNA-positive events.

### NSM or ASM Inhibition Differentially Impacts the EV Proteome

The distinctive features of EVs released by SMase inhibitor-treated cells prompted us to characterize their molecular cargoes. For this purpose, we first conducted liquid chromatography tandem mass spectrometry (LC-MS/MS) on triplicate cellular and EV specimens, following vehicle or inhibitor treatments, to define their proteomic signatures. Quantification of the total protein yield from each specimen type revealed no significant differences between cellular samples (**Figure 2A**). However, at the EV level, both NSM- and ASM-inhibited samples respectively exhibited trends of reduced and increased total protein compared to DMSO controls (**Figure 2B**). Multidimensional scaling (MDS) and Pearson correlation analyses demonstrated a high degree of similarity among all cellular protein specimens, while EV samples formed distinct treatment-defined clusters by MDS (**Figure 2C-D**). Differential expression analysis identified only a few proteins with varying expression in cellular specimens (**Figure S2A-D**). By contrast, SMase inhibitors had a more robust impact on the EV proteome composition relative to controls, with FTY720 EVs showing an increased expression of 451 and a decrease of 81 unique proteins, while GW4869 EVs exhibited an increase of only 22, but a marked reduction in 460 unique proteins (**Figure S2D**).

**FIGURE 2:**
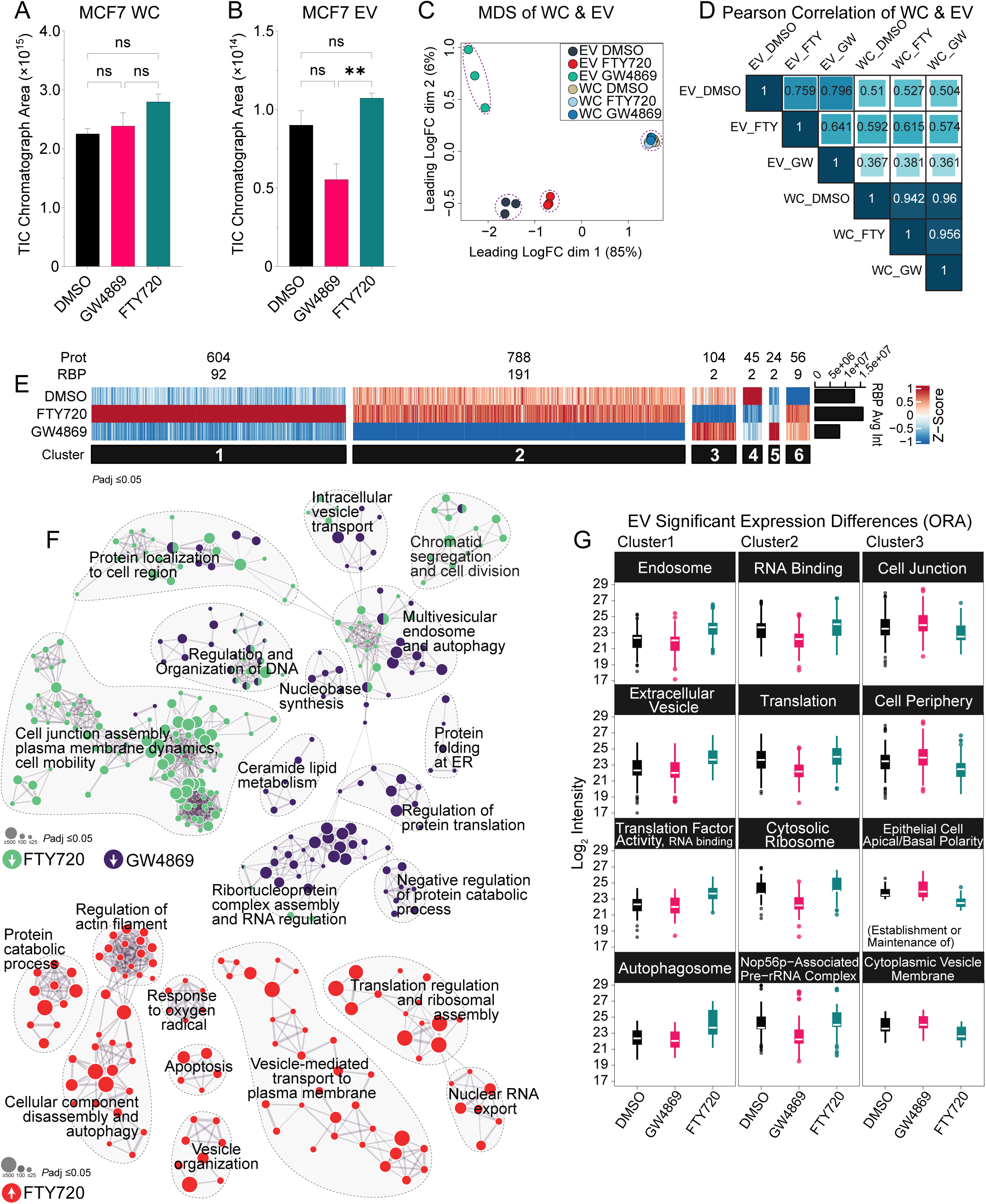
NSM or ASM Inhibition Differentially Impacts the EV Proteome. **A.** Total ion current (TIC) chromatograph area of proteins isolated from either control or SMase inhibitor-treated cells, as estimated by initial injection volume and total sample volume. **B.** TIC chromatograph area of proteins isolated from EVs derived from either control or SMase inhibitor-treated cells, as estimated by initial injection volume and total sample volume. **C.** Multidimensional scaling (MDS) of liquid chromatography tandem mass spectrometry (LC-MS/MS) proteomics data derived from cellular and EV purified proteins under control or SMase inhibitor-treated conditions. **D.** Pearson correlations of cellular and EV proteomes under control or SMase inhibitor-treated conditions. **E.** Clustered heatmap analysis of peptide intensities (relative Z-score) of EV proteomes from control or SMase inhibitor-treated conditions. Blue lines denote lower association, while red lines denote higher association of each protein relative to the other two groups. Only statistically significant proteins were included (adjusted p-value ≤ 0.05 and Log_2_ fold change ≤ -1 or ≥ 1). **F.** Cytoscape ClueGO analysis displaying a gene ontology (GO) biological process network of associated protein clusters, as identified by ORA, of statistically significant downregulated or upregulated EV proteins (adjusted p-value ≤ 0.05 and Log_2_ fold change ≤ -1 or ≥ 1) from control or SMase inhibitor-treated conditions. Green nodes represent downregulated FTY720 EVs, purple nodes represent downregulated GW4869 EVs, and red nodes represent upregulated FTY720 EVs. **G.** Boxplots of peptide intensities of EV protein clusters (subfigure E), categorized by overrepresentation analyses (ORA) with g:Profiler, displaying the total Log_2_ peptide intensities of all proteins in the groups and their changes under control or SMase inhibitor-treated conditions. Only statistically significant proteins were included (adjusted p-value ≤ 0.05 and Log_2_ fold change ≤ -1 or ≥ 1). Examples from clusters 1-3 displayed. Statistical analysis details for subfigures C-G are provided in the methodology. For subfigures A-B, the error bars represent the standard error of the mean (SEM) and the statistics were calculated by ordinary one-way ANOVA with Tukey post-hoc test (where ns=*P*>0.05, *=*P*≤0.05, **=*P*≤0.01, ***=*P*≤0.001, and ****= *P*≤0.0001) on GraphPad Prism.

To further analyze these differences at the EV level, we conducted clustering analysis on proteins exhibiting significant expression changes (Log_2_ fold change ≥ 1 or ≤ -1, adjusted p-value ≤ 0.05) in at least one of the comparison groups (DMSO vs GW4869 or FTY720). This analysis revealed six clusters of differential protein expression encompassing a total of 1,621 proteins (**Figure 2E**). Notably, the most predominant protein clusters (Clusters 1-3) exhibited opposite expression changes in response to drug treatments, highlighting the distinctive roles of NSM and ASM in influencing EV protein composition (**Figure 2E**). Strikingly, Clusters 1-2 contain a collection of 283 RNA-binding proteins (RBPs) exhibiting a significant depletion from EVs following NSM inhibition, while also displaying an increase in expression in ASM-inhibited EVs (**Figure 2E**, **S2E**).

Over-representation analysis (ORA) of EV protein clusters using g:Profiler was conducted to further contrast their functional signatures categorized by gene ontology (GO) terms. Network analysis of protein clusters, based on their enriched GO terms related to biological function, uncovered distinctive classes of proteins affected by each treatment (**Figure 2F**). For example, proteins specifically downregulated in the ASM inhibitor EV proteome (Clusters 3-4 in **Figure 2E**) were linked to cell junction assembly, plasma membrane dynamics, and cell division regulation (**Figure 2F**, green nodes). These same EVs showed an enrichment for proteins associated with protein catabolism, actin regulation, vesicle organization, and further included a substantial proportion of RBPs involved in diverse RNA regulatory functions, such as RNA nuclear export, translation regulation, and spliceosomal activity (**Figure 2F**, red nodes). In contrast, the NSM-inhibited EV proteome was depleted for proteins associated with autophagy, lipid metabolism, protein folding, and various RNA regulatory processes (**Figure 2F**, purple nodes). Quantifications of the total Log_2_ peptide intensities of all significant proteins within specific GO groups confirmed opposing effects in response to SMase inhibitors, denoting specificities related to intracellular compartments and biological function (**Figure 2G**, **S2F**). Most intriguingly, RBP content in EVs appeared to be largely influenced by NSM activity.

### Inhibition of ASM Alters the Transcript Diversity of EVs

Given the contrasting effect of SMase inhibitors on the EV targeting of RBPs, we next sought to assess their impact on EV RNA cargoes. RNA samples were extracted from control and SMase inhibitor-treated cellular and EV specimens. Quantification of total RNA content revealed minimal differences in cellular samples (**Figure 3A**); however, the total RNA content within EVs recovered from the drug treatments mirrored earlier measurements from nanoflow cytometry (**Figure 1J**), showing reduced recovery of RNA from GW4869 EVs and an increase in FTY720 EVs relative to control (**Figure 3B**). Sequencing on rRNA-depleted total RNA samples was performed using a NovaSeq platform, resulting in approximately 35 million paired-end reads per sample. Principal component analysis (PCA) (**Figure 3C**) and Pearson correlations (**Figure S3A**) of cellular and EV sequenced libraries revealed distinct class-specific distributions. While cellular samples exhibited highly correlated expression signatures, the transcriptomes of ASM-inhibited (FTY720) EVs were divergent compared to control or NSM-inhibited (GW4869) specimens (**Figure 3C**, **S3A**). This was reflected by differences in the proportion of reads obtained for different RNA biotypes, with mRNA and miscellaneous RNA (miscRNA) reads showing an increase in FTY720 EV specimens (**Figure 3D-E**), while reads for long non-coding RNAs (lncRNAs) and pseudogene transcripts were reduced (**Figure 3D-E**).

**FIGURE 3:**
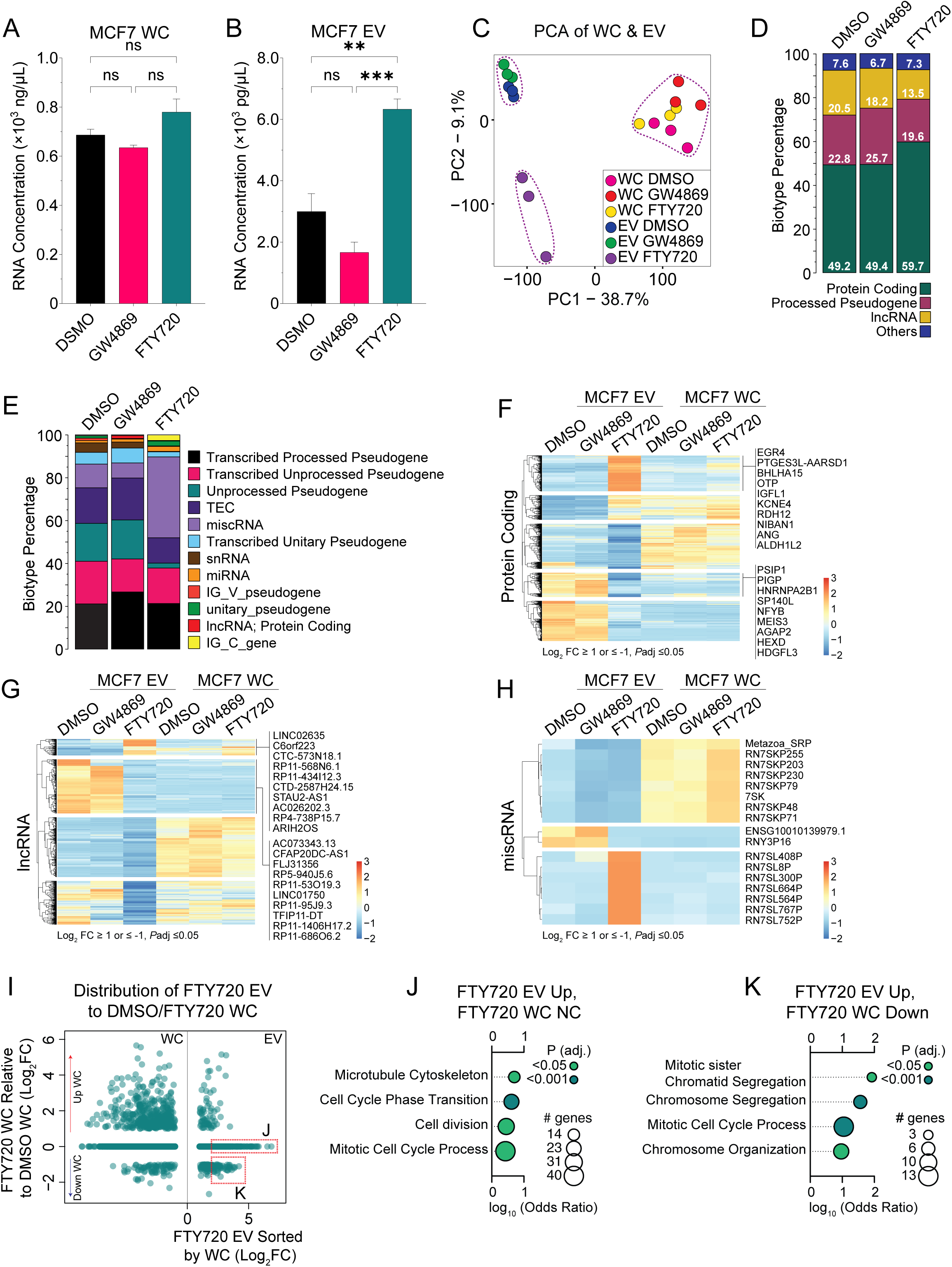
Inhibition of ASM Alters the Transcript Diversity of EVs. **A.** Spectrophotometer quantification of nucleic acid concentration, in nanograms per microliter (ng/uL), of total RNA isolated from either control or SMase inhibitor-treated cells. **B.** Spectrophotometer quantification of nucleic acid concentration, in picograms per microliter (pg/uL), of total RNA isolated from either control or SMase inhibitor-treated cells. **C.** Principal Component Analysis (PCA) of RNA sequencing data derived from cellular and EV purified RNAs from control or SMase inhibitor-treated specimens. **D.** Percentage distribution of RNA biotypes in EV transcriptomes from either control or SMase inhibitor-treated conditions. ’Others’ describe a varied group of minor transcripts, including but not limited to miscRNA, snRNA, miRNA, etc. (see Figure 4E). **E.** Percentage distribution of RNA biotypes in EV transcriptomes from either control or SMase inhibitor-treated conditions. This subfigure represents the ‘Others’ group in the Figure 4D. **F-H.** Clustered heatmap analyses of transcript biotypes (subfigures F=protein coding, G=lncRNA, H=miscRNA) expression (relative Z-score) in transcriptomes from control or SMase inhibitor-treated conditions. Blue lines denote lower expression, while red lines denote higher expression of all transcripts detected relative to the other two conditions. Genes displayed represent the 10 most differentially expressed transcripts from the highlighted clusters. Only statistically significant transcripts were included (adjusted p-value ≤ 0.05 and Log_2_ fold change ≤ -1 or ≥ 1). **I.** Dot plot illustrating the asymmetrical distribution of transcripts detected in acid sphingomyelinase (ASM)-inhibited EVs (FTY720) compared to ASM-inhibited cellular transcriptomes, normalized to control cellular transcriptomes. The upper dotted red box, labeled J, represents statistically significant upregulated transcripts (adjusted p-value ≤ 0.01 and log_2_ fold change ≥ 2) in ASM-inhibited EVs that remain unchanged in cells in response to the drug condition. The lower dotted red box, labeled K, represents statistically significant upregulated transcripts (adjusted p-value ≤ 0.01 and Log_2_ fold change ≥ 2) in ASM-inhibited EVs, which are simultaneously downregulated in ASM-inhibited cells (adjusted p-value ≤ 0.01 and Log_2_ fold change ≥ 2) in response to the drug condition. **J.** FuncAssociate v3.0 gene ontology (GO) enrichment analysis of statistically significant upregulated transcripts (adjusted p-value ≤ 0.01 and Log_2_ fold change ≥ 2) in ASM-inhibited EVs that remain unchanged in cells in response to the drug condition (see subfigure I). The calculated logarithm (base 10) of the odds (LOD) ratio and adjusted p-value are displayed. **K.** FuncAssociate v3.0 gene ontology (GO) enrichment analysis of statistically significant upregulated transcripts (adjusted p-value ≤ 0.01 and Log_2_ fold change ≥ 2) in ASM-inhibited EVs, which are simultaneously downregulated in ASM-inhibited cells (adjusted p-value ≤ 0.01 and log_2_ fold change ≥ 2) in response to the drug condition (see subfigure I). The calculated logarithm (base 10) of the odds (LOD) ratio and adjusted p-value are displayed. For subfigures A-B, the error bars represent the standard error of the mean (SEM) and the statistics were calculated by ordinary one-way ANOVA with Tukey post-hoc test (where ns=*P*>0.05, *=*P*≤0.05, **=*P*≤0.01, ***=*P*≤0.001, and ****= *P*≤0.0001) on GraphPad Prism.

Differential gene expression analysis also revealed that FTY720 EVs exhibit the most pronounced deviation from other conditions, characterized by the respective increased or decreased expression (Log_2_ fold change ≥ 1 or ≤ -1, adjusted p-value ≤ 0.05) of 820 or 2140 transcripts compared to DMSO treatment (**Figure S3B-E**) across various RNA biotypes (**Figure 3F-H**, **S3F-G**). Strikingly, RN7SL family transcripts were strongly enriched in FTY720 EVs (**Figure 3H**), while RN7SL1 was among the most prominently depleted transcripts in GW4869 EV specimens (**Figure S3F**). Further comparison of FTY720-EV (**Figure 3I**) or GW4869-EV (**Figure S3H**) samples to the normalized cellular expression data identified three groups of EV-detected RNAs: those upregulated in both cells and EVs, those upregulated in EVs but not altered in cells, and those upregulated in EVs but downregulated in cells (**Figure 3I**) in response to treatments. FTY720 EV-enriched mRNAs exhibit GO signatures for cell cycle regulatory processes, cell division, and microtubule cytoskeleton (**Figure 3I-K**), while no specific GO enrichments were observed for GW4869 EV-enriched mRNAs (**Figure S3H**). These results suggest that the sorting of RNA cargoes during EV biogenesis is nuanced, with a substantial influence from sphingomyelinase activity at these sites.

### RBPs as Major Differentially Regulated EV Cargoes in Response to NSM or ASM Inhibition

RBPs play pivotal roles as cellular regulators, modulating all aspects of RNA metabolism. Intriguingly, we generally observed opposing effects of SMase inhibitors on RBP levels within EVs. Indeed, while NSM inhibition with GW4869 resulted in the downregulation of a large number of detected RBPs (n=159; 43.7% of detected RBPs) compared to control EVs (**Figure 4A**), an opposite trend was observed with FTY720 treatment, which resulted in an enrichment of many RBPs (n=86; 23.6% of detected RBPs) in EV specimens (**Figure 4B**). Strikingly, a comparison of GW4869 versus FTY720 EV samples revealed an even stronger RBP enrichment signature (n=264; 72.5% of detected RBPs) in FTY720 specimens (**Figure 4C**). Considering these findings, we next investigated the effects of SMase inhibition on individual components of bipartite ribonucleoprotein (RNP) modules by comparing our proteomics and transcriptomic datasets.

**FIGURE 4:**
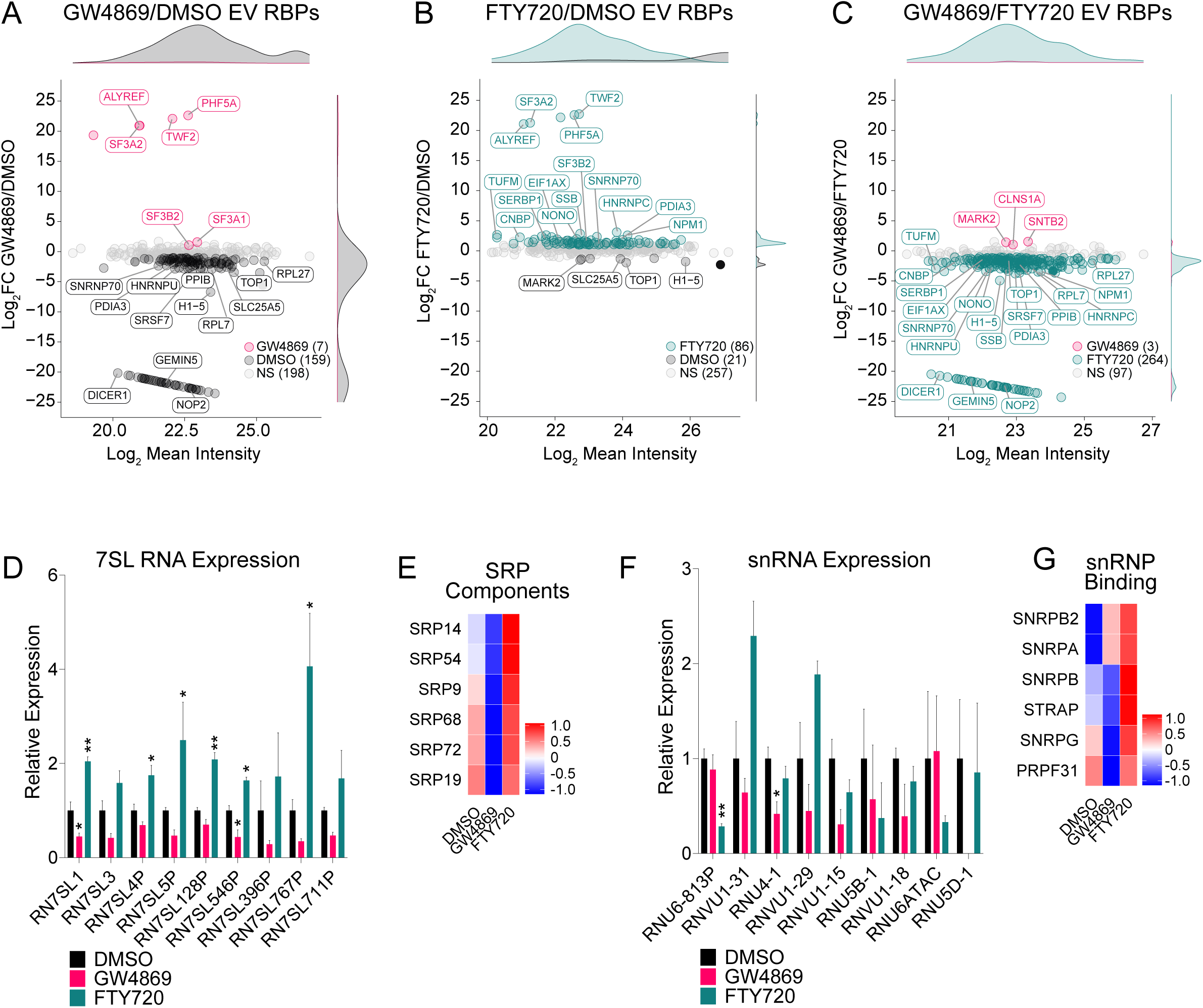
RBPs as Major Differentially Regulated EV Cargoes in Response to NSM or ASM Inhibition. **A.** MA plot of RNA-binding proteins (RBPs), detected by liquid chromatography tandem mass spectrometry (LC-MS/MS), illustrating the Log_2_ fold change and Log_2_ mean intensity of RBPs in EVs in response to neutral sphingomyelinase (NSM) inhibitor treatments (GW4869), relative to control EVs. **B.** MA plot of RNA-binding proteins (RBPs), detected by LC-MS/MS, illustrating the Log_2_ fold change and Log_2_ mean intensity of RBPs in EVs in response to acid sphingomyelinase (ASM) inhibitor treatments (FTY720), relative to control EVs. **C.** MA plot of RNA-binding proteins (RBPs), detected by LC-MS/MS, illustrating the Log_2_ fold change and Log_2_ mean intensity of RBPs in EVs in response to sphingomyelinase inhibitor treatments (GW4869/FTY720), relative to each other. **D.** Bar graphs illustrating the relative expression of 7SL RNAs (from Transcripts Per Million - TPMs), as detected by RNA sequencing (RNA-seq). The graphs display changes in transcript expression in EVs under both control and SMase inhibitor treatment conditions. **E.** Clustered heatmap analysis of peptide intensities (relative Z-score) depicting the changes in the expression of proteins associated with the signal recognition particle (SRP) within EVs under both control and SMase inhibitor-treated conditions. The color scale represents the strength of association, where blue signifies lower association and red indicates higher association for each protein compared to the other two groups. Only statistically significant proteins were included (adjusted p-value of ≤ 0.05 and a Log_2_ fold change of ≤ -1 or ≥ 1). **F.** Bar graphs illustrating the relative expression of U snRNAs (from TPMs), as detected by RNA-seq. The graphs display changes in transcript expression in EVs under both control and SMase inhibitor treatment conditions. **G.** Clustered heatmap analysis of peptide intensities (relative Z-score) depicting the changes in the expression of proteins associated with small nuclear ribonucleoproteins (snRNPs) of the spliceosome within EVs under both control and SMase inhibitor-treated conditions. The color scale represents the strength of association, where blue signifies lower association and red indicates higher association for each protein compared to the other two groups. Only statistically significant proteins were included (adjusted p-value of ≤ 0.05 and a Log_2_ fold change of ≤ -1 or ≥ 1). For subfigures D and F, the error bars represent the standard error of the mean (SEM) and the statistics were calculated by ordinary one-way ANOVA with Tukey post-hoc test (where ns=*P*>0.05, *=*P*≤0.05, **=*P*≤0.01, ***=*P*≤0.001, and ****= *P*≤0.0001) on GraphPad Prism.

While the impact of NSM inhibition on the EV RNA repertoire was relatively limited, specific examples emerged. Notably, the secretion of various members of the 7SL RNA family (e.g., RN7SL1, RN7SL3, and various 7SL pseudogenes), integral RNA components of the Signal Recognition Particle (SRP), exhibited significantly decreased expression in NSM inhibitor EVs but increased expression in ASM inhibitor EVs (**Figure 4D**). Moreover, these expression alterations were predominantly confined to EVs, with cellular transcripts remaining similar (**Figure S4A**). Intriguingly, the behavior of SRP proteins mirrored that of their RNA counterparts, displaying substantial depletion in NSM-inhibited EVs and more pronounced expression in ASM-inhibited EVs (**Figure 4E**). Another compelling example involves small nuclear ribonucleoproteins (snRNPs) associated with spliceosomal activity. Examination of the EV transcriptomic data revealed a trend of reduced expression of spliceosomal small nuclear RNAs (snRNAs) in NSM-inhibited EVs (**Figure 4F**). However, except for a few instances like RNVU1-31 and RNVU1-29, these RNAs did not exhibit an increase in ASM-inhibited EVs (**Figure 4F**). As before, these variations in expression were generally not observed in the cellular transcriptome samples (**Figure S4B**). Notably, this distinction was also evident at the proteomic level, where the secretion of several spliceosomal RBPs was downregulated in response to GW4869, while they were robustly increased in FTY720 samples (**Figure 4G**). Finally, several mRNAs associated with GO-defined ‘negative regulation of translation’, such as SAMD4A, AGO1, and NCL, were downregulated in the transcriptomes of ASM-inhibited EVs but remained relatively unaffected in NSM-inhibited EVs (**Figure S4C**). Importantly, the expression of these transcripts was generally consistent across cellular specimens (**Figure S4D**). Collectively, these findings indicate that while RNA and RBPs can be independently sorted into SMase-dependent EVs, there is also coherence in the secretion behavior of RNA and protein components of RNP modules.

### ASM-Inhibited EVs Enhance Protein Translation in Recipient Cells

Given the differential impact of SMase inhibitors on EV cargoes, we sought to investigate whether these treatments influenced the impact that these EVs might exert when transferred onto recipient cells. To address this, we established a co-culture system enabling us to quantify the phenotypic response of the recipient cell line, human MCF10A, to EVs isolated from control or inhibitor-treated MCF7 cells. Beginning with an equal number of cells, we obtained volumetrically equivalent samples of size exclusion chromatography (SEC)-purified EVs in 1x PBS from MCF7 cells. The quantity of proteins within the samples was assessed before loading onto recipient MCF10A cells (**Figure 5A**), revealing minimal differences in the total protein content. Additionally, a volumetric equivalent of 1x PBS was employed as a negative control.

**FIGURE 5:**
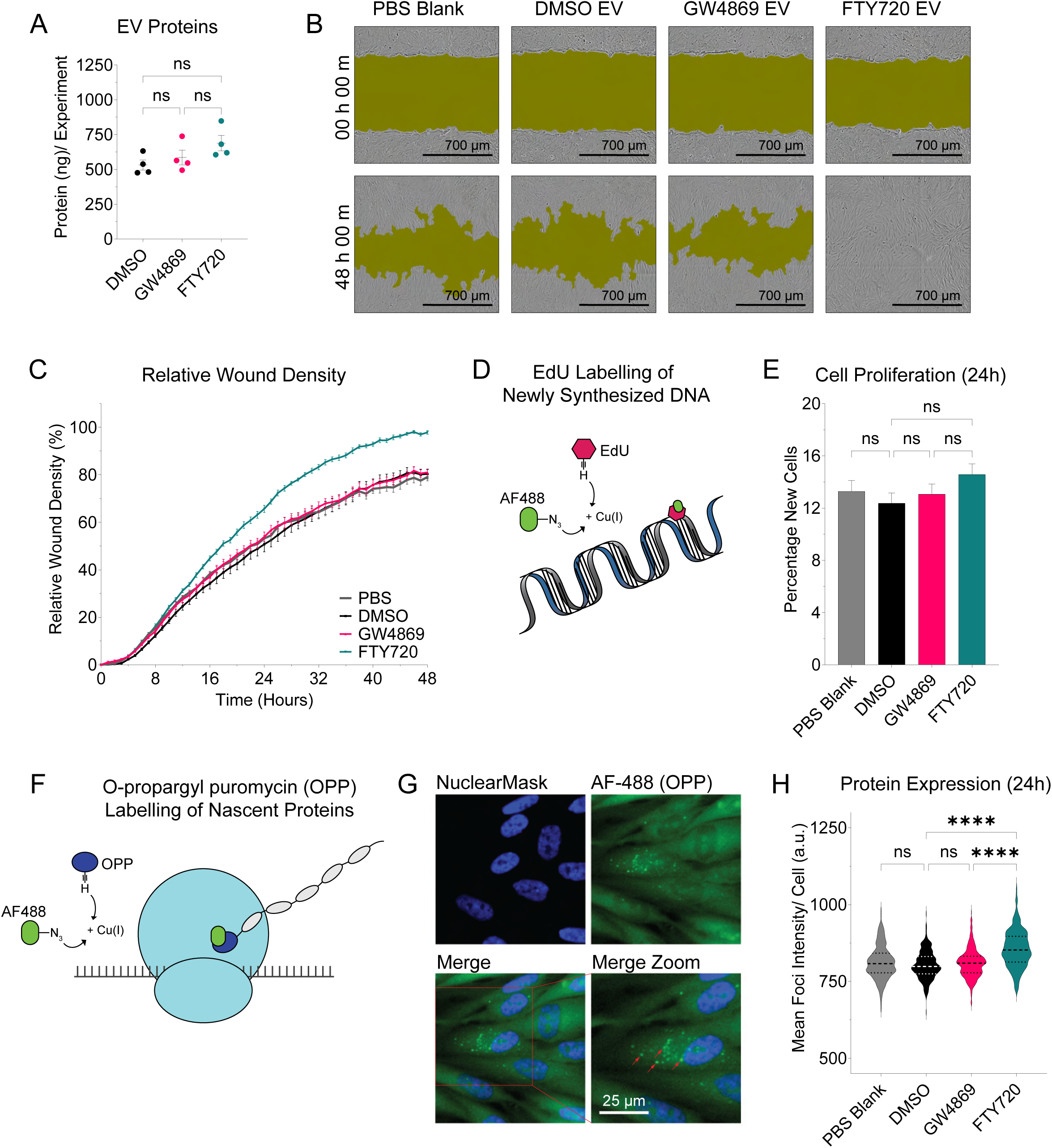
ASM-Inhibited EVs Enhance Protein Translation in Recipient Cells. **A.** Quantification of total EV proteins within the equal volumes of EVs utilized for co-culture experiments. The EVs used are present within equivalent isolation volumes purified from similar cell numbers that have been incubated under both control and SMase inhibitor conditions. **B.** Representative light microscope images of wounded MCF10A cells incubated with MCF7 EVs, obtained under control or SMase inhibitor conditions, as captured using the Incucyte Live-Cell Analysis System. The images were taken at the initiation (0h 00m) and conclusion (48h 00m) of the co-culture experiment. The Incucyte software applied a wound mask. The scale bar represents 700 μm. **C.** Quantification of the percentage Relative Wound Density (%RWD) in wounded MCF10A cells over a 48- hour period. The cells were incubated with MCF7 EVs obtained under control or SMase inhibitor conditions. Images were captured by an automated camera using the Incucyte Live-Cell Analysis System, and %RWD was calculated using its Scratch Wound Analysis Software Module. %RWD measures the spatial cell density in the wound area relative to the spatial cell density outside of the wound area at each time point. **D.** Schematic of the cell proliferation assay through the incorporation of 5-ethynyl-2’-deoxyuridine (EdU) in newly synthesized DNA, followed by the chemoselective ligation of Alexa Fluor™ 488 picolyl azide by a click chemistry reaction. **E.** Percentage of MCF10A cells in the total population labeled by 5-ethynyl-2’-deoxyuridine (EdU) after a 24-hour incubation with EdU and MCF7 EVs obtained under control or SMase inhibitor conditions. EdU incorporates into newly synthesized DNA, and quantification is based on high-content microscopy imaging. **F.** Schematic of the protein translation assay through the puromycylation of nascent proteins at the ribosome with O-propargyl puromycin (OPP), followed by the chemoselective ligation of Alexa Fluor™ 488 picolyl azide by a click chemistry reaction. **G.** Representative high-content screening microscopy images displaying nuclear stain (NuclearMask) and OPP-labelled protein foci (Alexa Fluor (AF)-488). Red arrows point to example OPP-labelled foci. **H.** Distribution of quantified mean intensities (arbitrary units – a.u.) of O-propargyl puromycin (OPP)-labeled foci in MCF10A cells following a 24-hour co-culture with MCF7 EVs obtained under control or SMase inhibitor conditions. OPP integrates into the nascent peptides of newly translated proteins, and quantification is performed based on images captured with high-content screening microscopy. For all graphs, unless otherwise stated, the error bars represent the standard error of the mean (SEM) and the statistics were calculated by ordinary one-way ANOVA with Tukey post-hoc test (where ns=*P*>0.05, *=*P*≤0.05, **=*P*≤0.01, ***=*P*≤0.001, and ****= *P*≤0.0001) on GraphPad Prism.

We first evaluated the impact of EV treatment on the migration properties of MCF10A cells in the context of a scratch wound-healing assay. Employing an automated live cell imaging approach, images of the wounded region were captured every hour over a 48-hour period (**Figure 5B-C**). The percentage relative wound density (%RWD), which measures the spatial cell density in the wound area relative to the spatial cell density outside of the wound area, was calculated at each time point. Strikingly, EVs from ASM-inhibited conditions exhibited a higher wound closure rate, fully populating the wounded area within 48 hours (**Figure 5B**). Indeed, analysis of the %RWD revealed a statistically significant increase in wound density in MCF10A wounded specimens co-cultured with EVs purified from FTY720-treated cells starting at 16 hours, persisting until 48 hours, while no discernible effects were observed across the other conditions (**Figure 5C**).

Considering the observed effects of inhibitor treatments on mRNAs associated with cell division, we first employed 5-ethynyl-2’-deoxyuridine (EdU) to label newly synthesized DNA (**Figure 5D, S5A**) and quantify cell proliferation within the recipient MCF10A cell population. Quantification of the percentage of new MCF10A cells across different conditions revealed no difference between EV treatment specimens (**Figure 5E**, **S5A-C**), indicating that the observed enhancement in wound closure activity is not due to an increase in cell number. Guided by the cargo sorting effects on translation-associated machineries, such as SRP protein and 7SL RNA components, we next evaluated the impact of EVs on translational responses in recipient cells. Employing high-content screening microscopy, we quantified the intensity of O-propargyl-puromycin (OPP)-labeled foci, indicative of nascent puromycylated peptides (**Figure 5F-G**), in MCF10A cells co-cultured with MCF7 EV specimens. Our quantifications revealed that MCF10A cells co-cultured with ASM-inhibited EVs displayed an increased foci intensity signal starting at 6 hours (**Figure S5E**), and this effect persisted up to 24 hours (**Figure 5H**, **S5D-F**). In contrast, no discernible effects on MCF10A cell translational activity were observed with EVs from DMSO- or GW4869-treated MCF7 cells. We conclude that EV specimens derived from FTY720 treated cells have the capacity to enhance translation upon transfer to recipient cells, which likely explains the effects on cell migration.

## DISCUSSION

In recent years, extracellular vesicles (EVs) have been recognized as key players in intercellular communication; however, their complex landscape of heterogeneity that presents challenges in elucidating the distinct functions of subpopulations ^17,45^. EV biogenesis is primarily regulated by two mechanisms: the endosomal sorting complex required for transport (ESCRT)-dependent, and the ESCRT-independent mechanism, which is driven by sphingomyelinases (SMase) ^46,47^. Neutral (NSM) and acid (ASM) SMases, which convert sphingomyelin to ceramide, have been associated with different EV subpopulations, suggesting they play distinct roles in EV formation ^20–27^.

Various chemical inhibitors of SMases have been identified, offering an efficient approach for functional exploration compared to genetic knockdown methods ^48–55^. In this study, we inhibited NSM with GW4869 and ASM with FTY720 ^21,26,30–33^, allowing us to unveil their distinct contributions to the heterogenous EV population. Indeed, our initial observations revealed that neither GW4869 nor FTY720 could completely halt EV release (**Figure 1D-F**). This is a phenomenon that has been previously observed with chemical or genetic knockdowns of EV biogenesis components ^56–64^. However, each SMase inhibitor uniquely impacted the heterogeneous EV population, affecting their size distribution (**Figure 1E-H**), the quantity released (**Figure 1I**), and the composition of the cargoes they transported (**Figure 1J**).

Investigations into the proteomic composition of altered EV populations revealed a significant impact of sphingolipid metabolism on EV protein composition. While cellular protein expression remained consistent across inhibitor conditions (**Figure 2A**, **S2A-D**), EV proteins responded differently, showing changes in both relative quantities (**Figure 2B**) and contents (**Figure 2E-G**, **S2D-F**). Notably, distinct protein groups were affected uniquely by each treatment, often exhibiting opposing trends, such as a notable shift in the abundance of RNA-binding proteins (**Figure 2E**). Gene ontology (GO) and network analyses indicated that ASM inhibition led to the downregulation of proteins linked to plasma membrane dynamics, whereas NSM inhibition resulted in decreased levels of proteins associated with endosomal and autophagic activity (**Figure 2F-G**). These findings align with prior research on the role of NSM in exosome biogenesis ^21,22,65^ , highlighting its distinct mechanism compared to the ESCRT-dependent pathway ^30^. Interestingly, ASM inhibition also led to the downregulation of EV proteins linked to the multivesicular endosome (**Figure 2F**), potentially related to ASM’s functions in the lysosome and secretory autophagy ^66–68^, although further investigation is necessary.

Transcriptomic profiling uncovered a nuanced role for SMases in EV-mediated RNA release. Surprisingly, while NSM inhibition decreased extracellular RNA levels (**Figure 1J**, **3B**), GW4869-treated EVs showed only moderate transcriptomic changes (**Figure 3C-H**, **S3D-H**). In contrast, ASM inhibition increased extracellular RNA release but decreased it per vesicle (**Figure 1J**, **3B**). Additionally, ASM inhibition induced significant shifts in EV biotype composition, affecting mRNA, lncRNA, miscRNA, and pseudogene transcripts (**Figure 3D-H**, **S3G**). Notably, ASM-inhibited EVs exhibited enriched RN7SL family transcripts, while NSM inhibition depleted RN7SL1 in EVs. These results imply a complex RNA trafficking mechanism involving multiple sorting pathways, such as ESCRT ^69^.

The bioactive cargoes within EVs play critical roles in intercellular signaling, influencing the physiological state of recipient cells ^18^. Our data clearly demonstrated distinct alterations in the heterogeneous EV population, prompting an investigation into whether these changes could impact their effects on recipient cells. Co-culture experiments involving MCF7 SMase inhibitors EVs and recipient MCF10A cells revealed significant influences on cell behavior in response to FTY720 EVs (**Figure 5B-H**). Specifically, a scratch-wound healing assay showed accelerated migration of MCF10A cells when exposed to ASM-inhibited EVs (**Figure 5B-C**). Motivated by noteworthy findings in our transcriptomic and proteomic datasets, particularly regarding changes in the export of components associated with translational and cell proliferation machinery, we explored whether these mechanisms were altered in MCF10A cells, potentially elucidating the observed wound closure phenotypes. Interestingly, tracking of new cell proliferation using 5-ethynyl-2’-deoxyuridine (EdU) revealed no significant changes (**Figure 5E**); however, assessment of nascent peptide puromycylation with O-propargyl-puromycin indicated increased protein expression in MCF10A cells in response to FTY720 EVs (**Figure 5H**).

Here, we provide a comprehensive examination of the role of sphingolipid metabolism in shaping the cargo composition of EVs. The evidence presented strongly underscores the pivotal involvement of sphingomyelinases in EV biogenesis and molecular composition, revealing distinct contributions from NSM and ASM. We demonstrate that the generation of EVs is reliant on the specific enzyme variant engaged (NSM or ASM), and cells may resort to alternative pathways when one is inhibited. Furthermore, our data highlights the unique roles of NSM and ASM in cargo sorting, where NSM inhibition downregulates RNP modules, and ASM inhibition differentially impacts RNA export. Notably, NSM-generated EVs emerge as major conduits for RBP export. Intriguingly, alterations in the EV population due to SMase inhibition can affect their ability to facilitate intercellular communication. These insights suggest heightened significance for specific EV subpopulations in the context of intercellular communication and function.

## LIMITATIONS OF THE STUDY

While this study provides valuable insights into the roles of NSM and ASM in EV biogenesis, some limitations should be acknowledged. Although our sphingomyelinase activity assays confirm the expected effects of SMase inhibitors, conducting complementary genetic experiments would enhance the robustness of our findings. Additionally, the study focused solely on NSM and ASM pathways, without investigating other mechanisms such as ESCRT-dependent biogenesis, which also play crucial roles in EV formation. Therefore, changes in EV composition and function observed in response to NSM and ASM inhibition may not solely stem from these pathways, as contributions from alternative biogenesis pathways cannot be discounted. Furthermore, while the study identifies proteomic and transcriptomic changes in response to SMase inhibition, the functional implications for intercellular communication and disease require further elucidation.

## ACKNOWLEDGMENTS

The authors are grateful for the invaluable support from the Molecular Biology and Functional Genomics platform (Sarah Boissel and team), and the Mass Spectrometry and Proteomics platform (Denis Faubert and Josée Champagne), at the *Institut de Recherches Cliniques de Montréal* (IRCM). Special thanks also go to S. Kelly Sears, Jeannie Mui, and the Facility for Electron Microscopy Research of McGill University, for their assistance. Additionally, the authors extend their appreciation to Janusz Rak, Dongsic Choi, and Laura Montermini for their generous training and support throughout the project. This research was made possible by grants awarded to E.L. from the Canadian Institutes of Health Research (CIHR), the Natural Sciences and Engineering Research Council of Canada (NSERC), and the Fonds de Recherche du Québec—Santé (FRQS). E.L. is a distinguished resarch scholar of the FRQS. J.C.A.P. was a recipient of doctoral scholarships provided by the FRQS, McGill University, and the IRCM.

## AUTHOR CONTRIBUTIONS

Conceptualization, J.C.A.P. and E.L.; Methodology, J.C.A.P., E.K., and E.L.; Investigation, J.C.A.P., Y.C., E.S., and E.K.; Formal Analysis, J.C.A.P.; S.B., J.B., E.K., and E.L.; Data Curation, J.C.A.P., S.B., J.B., and E.S.; Visualization, J.C.A.P., S.B., J.B., and E.S.; Writing – Original Draft: J.C.A.P.; Writing – Review & Editing, J.C.A.P., S.B., J.B., Y.C., E.S., E.K., and E.L.; Project Administration, J.C.A.P. and E.L.; Funding Acquisition, E.L.; Resources, E.L.; Supervision, E.L.

## SUPPLEMENTARY FIGURE LEGENDS

**FIGURE S1:**
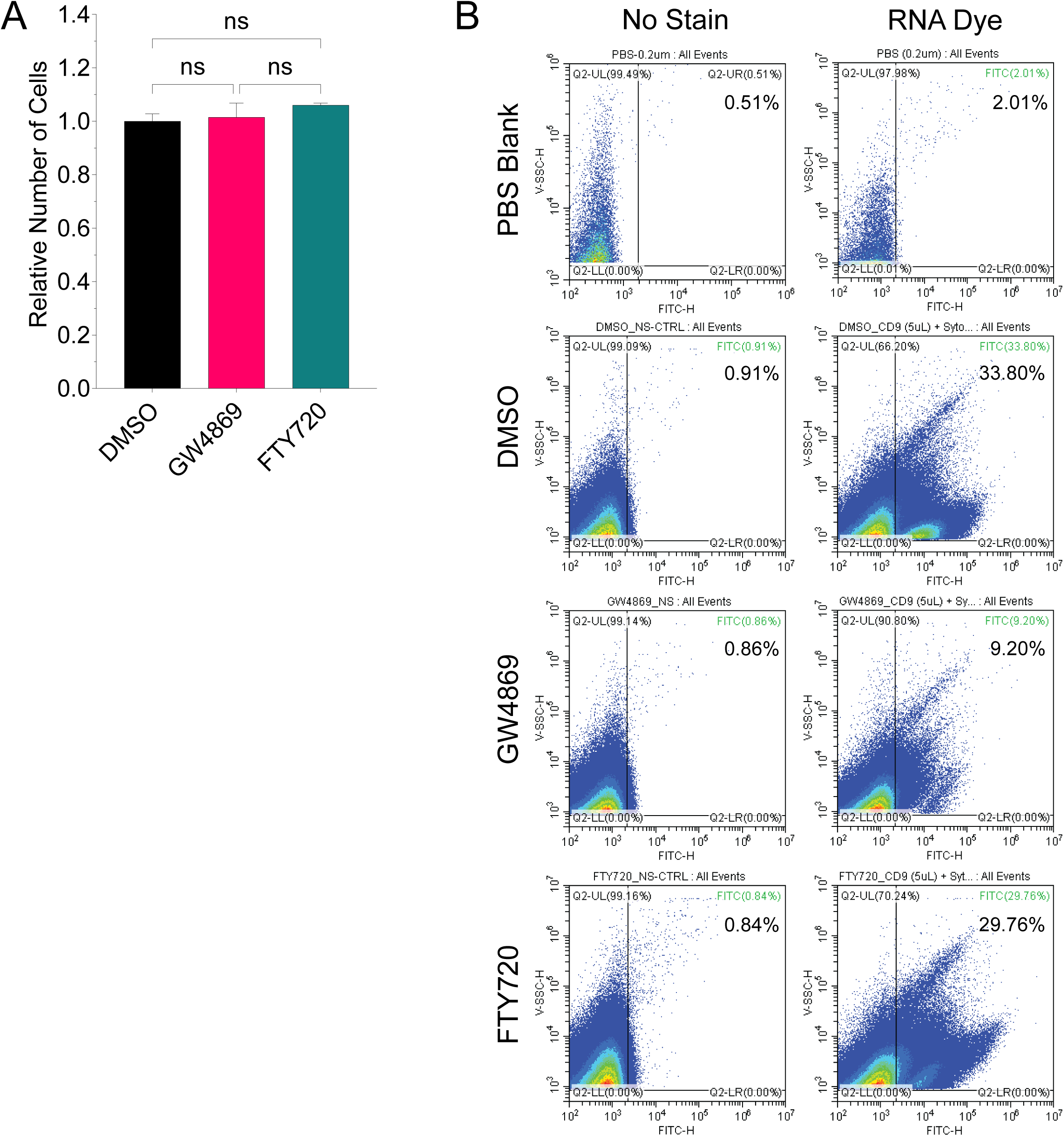
Sphingolipid Metabolism Influences the Release of EV RNA from Cells. **A.** Quantifications of the relative total number of cells conducted under control conditions (DMSO 0.2%) or sphingomyelinase (SMase) inhibitor conditions (10 μM GW4869 or 5 μM FTY720). The assessment was performed at the time of EV harvesting after 36 hours of incubation. **B.** Representative nanoflow cytometry profiles as captured by the CytoFLEX nanoflow cytometry platform. RNA-positive events were detected by staining of concentrated cell-conditioned media samples with SYTO RNASelect, followed by the purification of nanoparticles using size exclusion chromatography (SEC). Percent values represent the percentage of particles detected under FITC (RNA dye). For subfigure A, the error bars represent the standard error of the mean (SEM) and the statistics were calculated by ordinary one-way ANOVA with Tukey post-hoc test (where ns=*P*>0.05, *=*P*≤0.05, **=*P*≤0.01, ***=*P*≤0.001, and ****= *P*≤0.0001) on GraphPad Prism.

**FIGURE S2:**
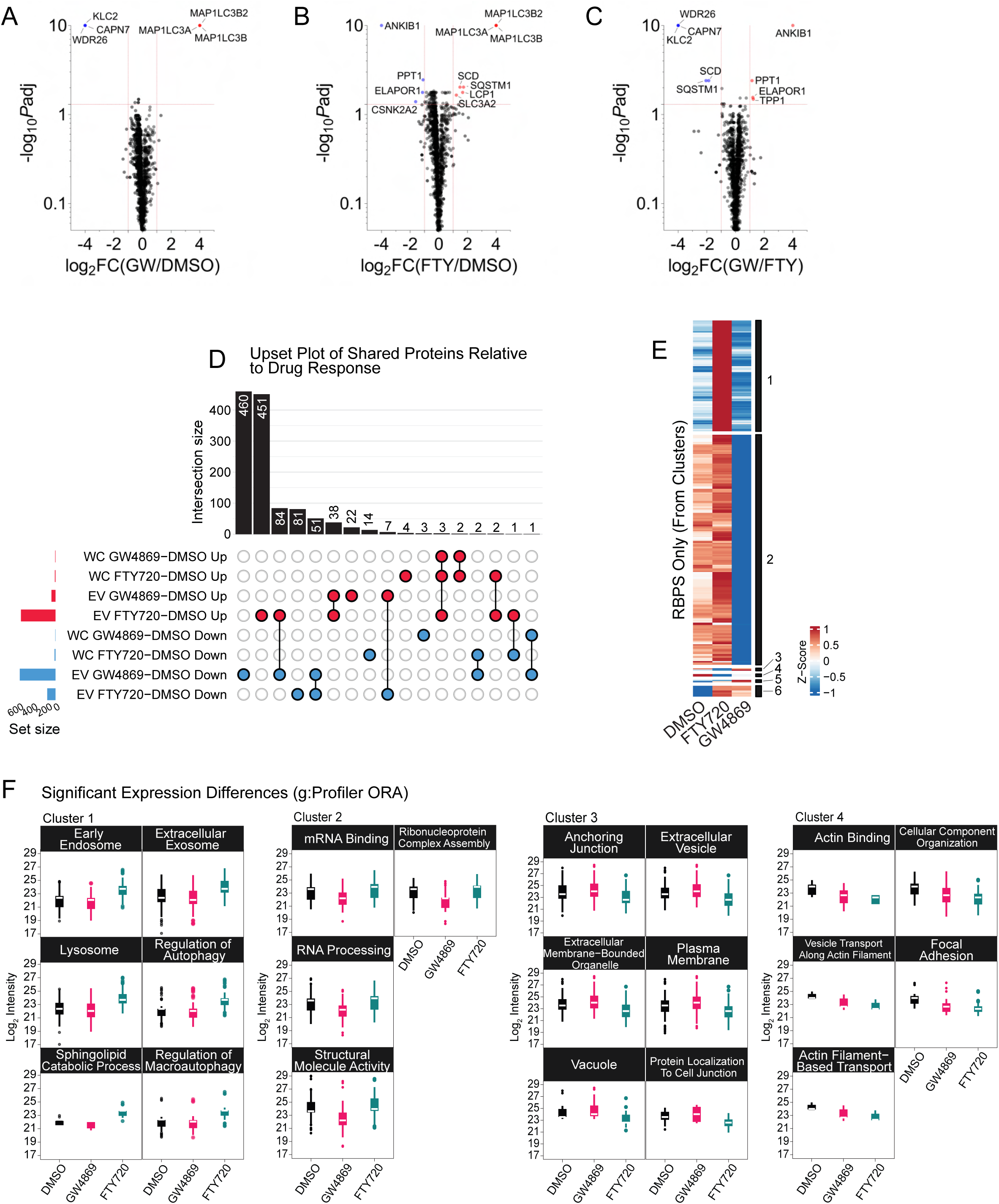
NSM or ASM Inhibition Differentially Impacts the EV Proteome. **A.** Volcano plot of cellular proteomes illustrating the Log_2_ fold change in protein expression in response to neutral sphingomyelinase (NSM) inhibitor treatments (GW=GW4869), relative to control cells. **B.** Volcano plot of cellular proteomes illustrating the Log_2_ fold change in protein expression in response to acid sphingomyelinase (ASM) inhibitor treatments (FTY=FTY720), relative to control cells. **C.** Volcano plot of cellular proteomes illustrating the Log_2_ fold change in protein expression in response to NSM inhibitor treatments (GW=GW4869), relative to ASM inhibitor treatments (FTY=FTY720). For subfigures A-C, the blue dots represent statistically significant downregulated transcripts (adjusted p-value ≤ 0.05, Log_2_ fold change ≤ -1), while the red dots represent statistically significant upregulated transcripts (adjusted p-value ≤ 0.05, Log_2_ fold change ≥ 1). **D.** UpSet plot of all LC-MS/MS-detected proteins in cellular and EV protein specimens derived from control or SMase inhibitor-treated conditions. The red nodes represent statistically significant upregulated transcripts (adjusted p-value ≤ 0.05, Log_2_ fold change ≥ 1), while the blue nodes represent statistically significant downregulated transcripts (adjusted p-value ≤ 0.05, Log_2_ fold change ≤ -1). The black nodes represent unregulated (unchanged) proteins. **E.** Clustered heatmap analysis of RNA-binding protein (RBP) peptide intensities (relative Z-score) of EV proteomes from control or SMase inhibitor-treated conditions. Blue lines denote lower association, while red lines denote higher association of individual RBPs relative to SMase inhibitor conditions. Only statistically significant RBPs were included (adjusted p-value ≤ 0.05 and Log_2_ fold change ≤ -1 or ≥ 1). **F**. Boxplots of peptide intensities of EV protein clusters (Figure 2E), categorized by overrepresentation analyses (ORA) with g:Profiler, displaying the total Log_2_ peptide intensities of all proteins in the groups and their changes under control or SMase inhibitor-treated conditions. Only statistically significant proteins were included (adjusted p-value ≤ 0.05 and Log_2_ fold change ≤ -1 or ≥ 1). Examples from clusters 1-4 displayed.

**FIGURE S3:**
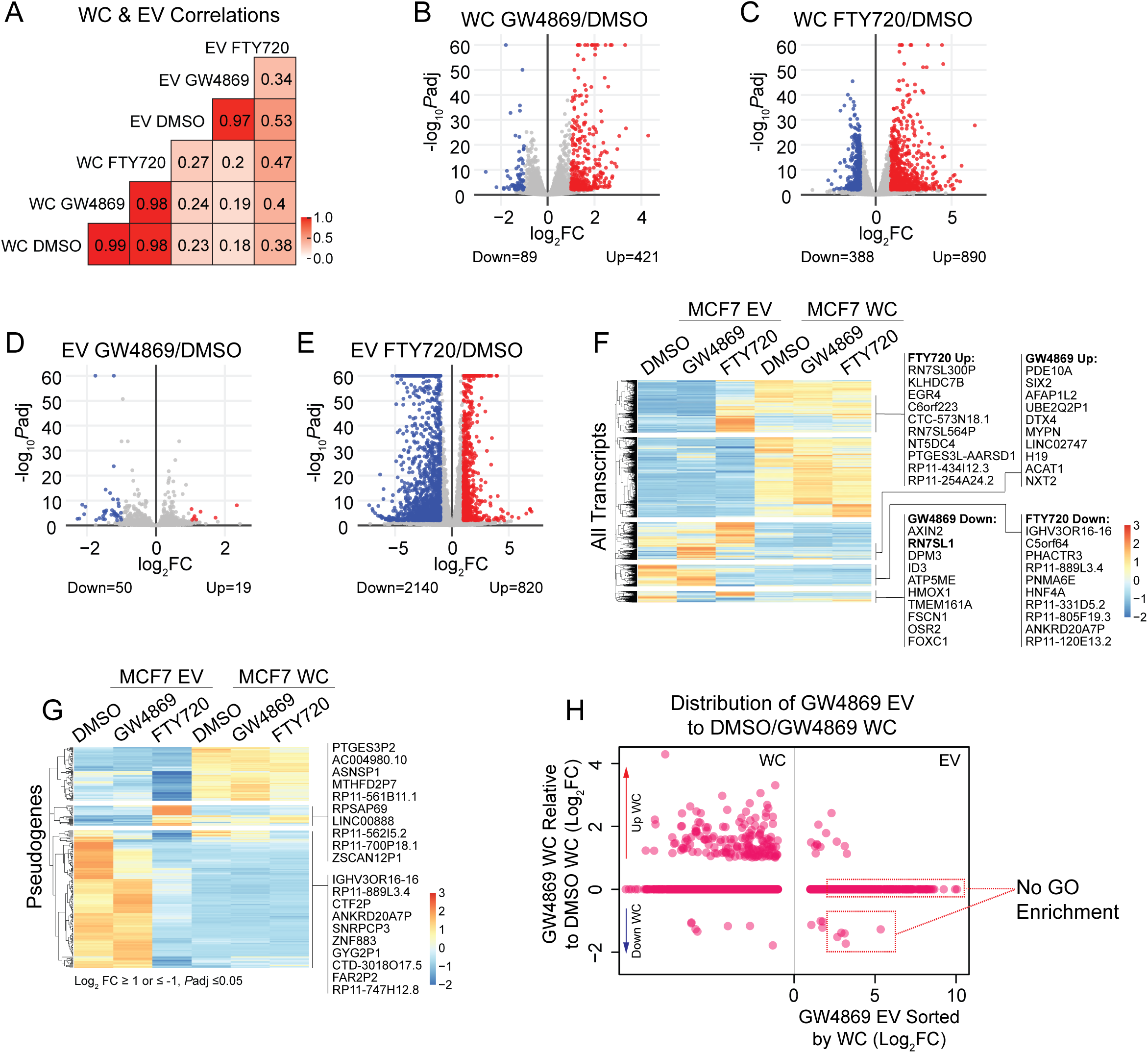
Inhibition of ASM Alters the Transcript Diversity of EVs. **A.** Pearson correlations of cellular and EV transcriptomes under control or SMase inhibitor-treated conditions. **B-E.** Volcano plot of cellular (subfigures D-E) and EV (subfigures F-G) transcriptomes illustrating the Log_2_ fold change in RNA expression in response to acid sphingomyelinase (ASM) (FTY720) or neutral sphingomyelinase (NSM) (GW4869) inhibitor treatments, relative to control conditions. The blue dots represent statistically significant downregulated transcripts (Log_2_ fold change ≤ -1, adjusted p-value ≤ 0.05). The red dots represent statistically significant upregulated transcripts (Log_2_ fold change ≥ 1, adjusted p-value ≤ 0.05). **F.** Clustered heatmap analysis of global differentially expressed transcripts (relative Z-score) in transcriptomes from control or SMase inhibitor-treated conditions. Blue lines denote lower expression, while red lines denote higher expression of all transcripts detected relative to the other two conditions. Genes displayed represent the 10 most differentially expressed transcripts from the highlighted clusters. Only statistically significant transcripts were included. **G.** Clustered heatmap analyses of pseudogene transcript expression (relative Z-score) in EV transcriptomes from control or SMase inhibitor-treated conditions. Blue lines denote lower expression, while red lines denote higher expression of all transcripts detected relative to the other two conditions. Genes displayed represent the 10 most differentially expressed transcripts from the highlighted clusters. Only statistically significant transcripts were included (adjusted p-value ≤ 0.05 and Log_2_ fold change ≤ -1 or ≥ 1). **H.** Dot plot illustrating the asymmetrical distribution of transcripts detected in neutral sphingomyelinase (NSM)-inhibited EVs (GW4869) compared to NSM-inhibited cellular transcriptomes, normalized to control cellular transcriptomes. The upper dotted red box represents statistically significant upregulated transcripts (Log_2_ fold change ≥ 2, adjusted p-value ≤ 0.01) in NSM-inhibited EVs that remain unchanged in cells in response to the drug condition. The lower dotted red box represents statistically significant upregulated transcripts (Log_2_ fold change ≥ 2, adjusted p-value ≤ 0.01) in NSM-inhibited EVs, which are simultaneously downregulated in NSM-inhibited cells (Log_2_ fold change ≤ -2, adjusted p-value ≤ 0.01) in response to the drug condition. FuncAssociate v3.0 gene ontology (GO) enrichment analysis of statistically significant upregulated transcripts was performed. No enrichment was detected with this tool.

**FIGURE S4:**
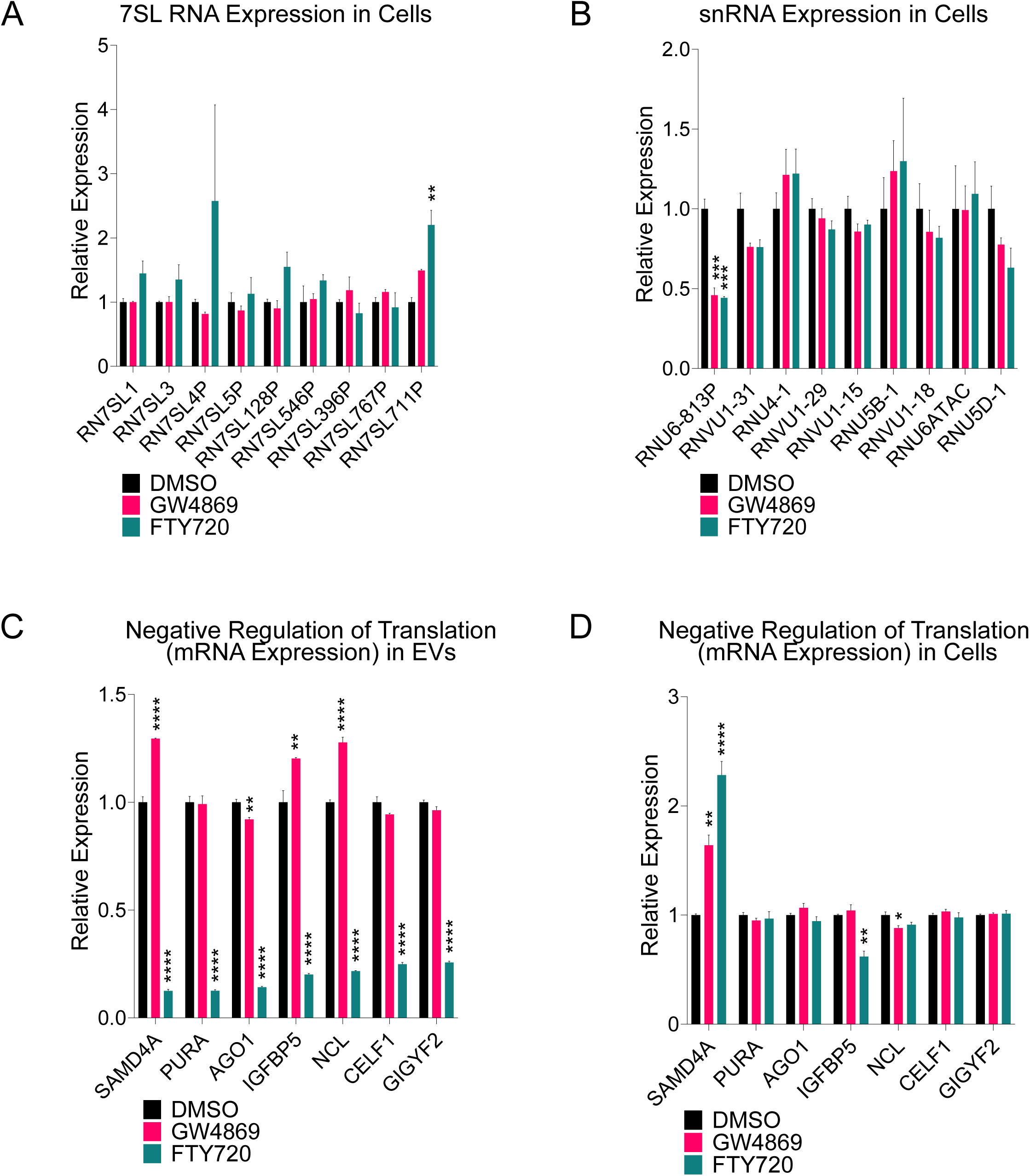
RBPs as Major Differentially Regulated EV Cargoes in Response to NSM or ASM Inhibition. **A.** Bar graphs illustrating the relative expression of 7SL RNAs (from Transcripts Per Million - TPMs), as detected by RNA sequencing (RNA-seq). The graphs display changes in transcript expression in cells under both control and SMase inhibitor treatment conditions. **B.** Bar graphs illustrating the relative expression of U snRNAs (from TPMs), as detected by RNA-seq. The graphs display changes in transcript expression in cells under both control and SMase inhibitor treatment conditions. **C.** Bar graphs illustrating the relative expression of mRNAs associated to the negative regulation of translation (from TPMs), as detected by RNA-seq. The graphs display changes in transcript expression in cells under both control and SMase inhibitor treatment conditions. **D.** Bar graphs illustrating the relative expression of mRNAs associated to the negative regulation of translation (from TPMs), as detected by RNA-seq. The graphs display changes in transcript expression in EVs under both control and SMase inhibitor treatment conditions. For all graphs, the error bars represent the standard error of the mean (SEM) and the statistics were calculated by ordinary one-way ANOVA with Tukey post-hoc test (where ns=*P*>0.05, *=*P*≤0.05, **=*P*≤0.01, ***=*P*≤0.001, and ****= *P*≤0.0001) on GraphPad Prism.

**FIGURE S5:**
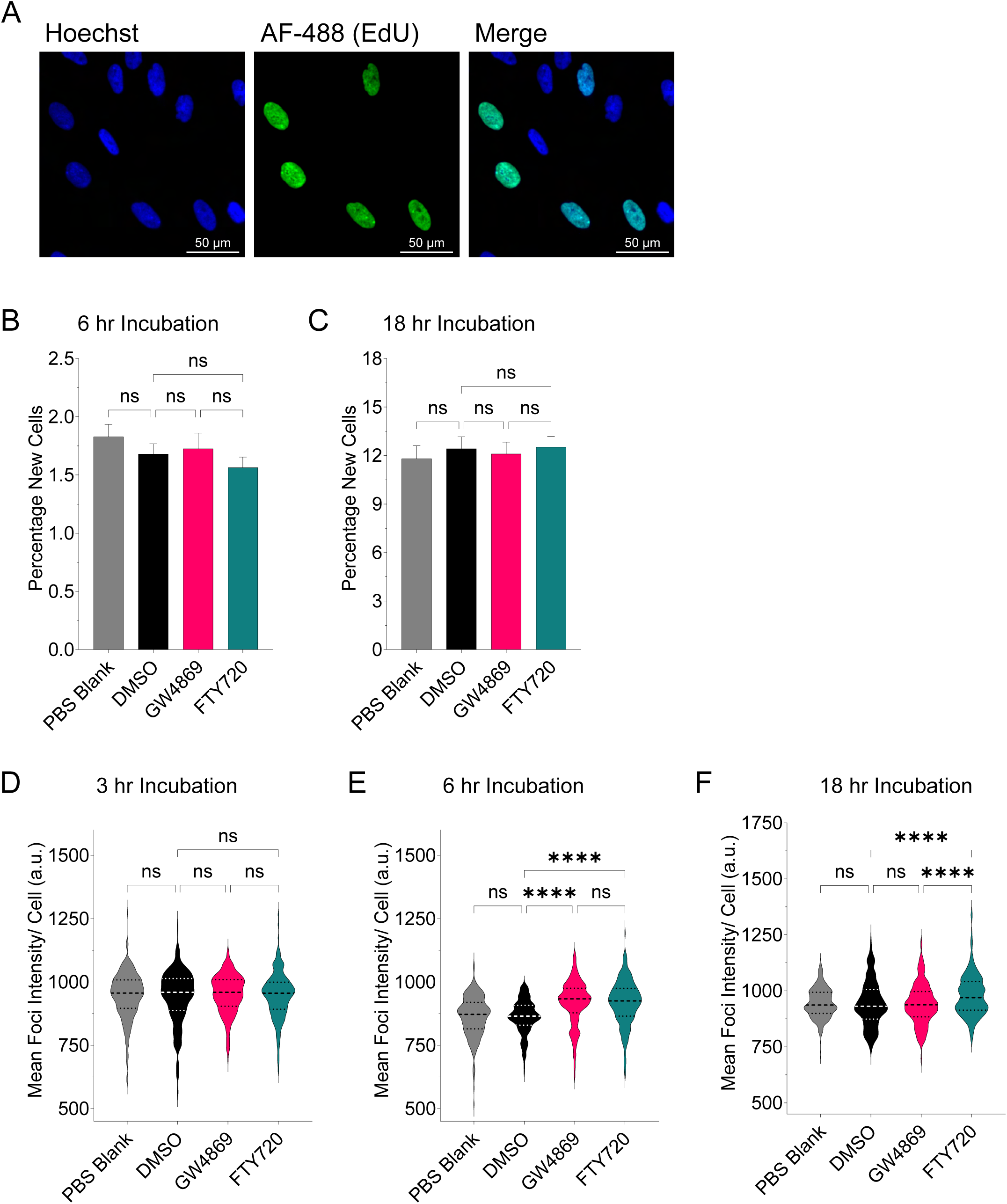
ASM-Inhibited EVs Enhance Protein Translation in Recipient Cells. **A.** Representative high-content screening microscopy images displaying nuclear stain (Hoechst) and new cells with incorporated EdU (Alexa Fluor (AF)-488). **B-C.** Percentage of MCF10A cells in the total population labeled by EdU after incubation with MCF7 EVs obtained under control or SMase inhibitor conditions. Quantification is based on high-content microscopy imaging on images captured over a time course: 6 hours (subfigure D), 18 hours (subfigure E), and 24 hours (Main Figure 6D). **D-F.** Distribution of quantified mean intensities (arbitrary units – a.u.) of OPP-labeled foci in MCF10A cells following co-culture with MCF7 EVs obtained under control or SMase inhibitor conditions. Quantification is performed on images captured with high-content screening over a time course: 3 hours (subfigure A), 6 hours (subfigure B), 18 hours (subfigure C), and 24 hours (Main Figure 6E). For all graphs, the error bars represent the standard error of the mean (SEM) and the statistics were calculated by ordinary one-way ANOVA with Tukey post-hoc test (where ns=*P*>0.05, *=*P*≤0.05, **=*P*≤0.01, ***=*P*≤0.001, and ****= *P*≤0.0001) on GraphPad Prism.

